# Data driven model discovery and interpretation for CAR T-cell killing using sparse identification and latent variables

**DOI:** 10.1101/2022.09.22.508748

**Authors:** Alexander B. Brummer, Agata Xella, Ryan Woodall, Vikram Adhikarla, Heyrim Cho, Margarita Gutova, Christine E. Brown, Russell C. Rockne

## Abstract

In the development of cell-based cancer therapies, quantitative mathematical models of cellular interactions are instrumental in understanding treatment efficacy. Efforts to validate and interpret mathematical models of cancer cell growth and death hinge first on proposing a precise mathematical model, then analyzing experimental data in the context of the chosen model. In this work, we present the first application of the sparse identification of non-linear dynamics (SINDy) algorithm to a real biological system in order discover cell-cell interaction dynamics in *in vitro* experimental data, using chimeric antigen receptor (CAR) T-cells and patient-derived glioblastoma cells. By combining the techniques of latent variable analysis and SINDy, we infer key aspects of the interaction dynamics of CAR T-cell populations and cancer. Importantly, we show how the model terms can be interpreted biologically in relation to different CAR T-cell functional responses, single or double CAR T-cell-cancer cell binding models, and density-dependent growth dynamics in either of the CAR T-cell or cancer cell populations. We show how this data-driven model-discovery based approach provides unique insight into CAR T-cell dynamics when compared to an established model-first approach. These results demonstrate the potential for SINDy to improve the implementation and efficacy of CAR T-cell therapy in the clinic through an improved understanding of CAR T-cell dynamics.

## INTRODUCTION

Dynamical systems modeling is one of the most successfully implemented methodologies throughout mathematical oncology (1). Applications of these *model first* approaches have lead to important insights in fundamental cancer biology as well as the planning and tracking of treatment response for patient cohorts (2, 3, 4, 5, 6, 7, 8, 9). Simultaneously, the last twenty years have seen explosive growth in the study and application of data-driven methods. These *data first* approaches, initially implemented as machine learning methods for imaging and genomics analyses, have seen much success (10, 11). However, such approaches are often limited to classification problems and fall short when the intention is to identify and validate mathematical models of the underlying dynamics. Recent efforts by us and others have aimed to develop methodologies that bridge these *model first* and *data first* approaches. (12, 13, 14).

In this work, we combine the methods of latent variable discovery and sparse identification of nonlinear dynamics (SINDy) (15, 16, 17) to analyze experimental *in vitro* cell killing assay data for chimeric antigen receptor (CAR) T-cells and glioblastoma cancer cells (18). This experimental data, featuring high temporal resolution, offers a unique opportunity to conduct an *in situ* test of the SINDy model discovery method. Interpretation of the discovered SINDy model is conducted under the expectation of a predator-prey interaction model in which the cancer cells function as the prey and the CAR T-cells the predator (19).

Predator-prey systems are a broad class of ordinary differential equations (ODEs) that aim to characterize changes in populations between two or more groups of organisms in which at least one survives via predation on another. Originally applied to the study of plant herbivory (20) and fishery monitoring (21) in the early 20^th^ century, predator-prey models have since become a workhorse of ecology, evolutionary biology, and most recently mathematical oncology (19, 22). Importantly, predator-prey models underpin much of the computational modeling of CAR T-cell killing, particularly in the context of *in vitro* cell killing assays (23, 7). An important example of these is the CAR T-cell Response in GliOma (CARRGO) model, a model that characterizes the *in vitro* interactions between CAR T-cells and glioma cells (18). The CARRGO model has shed light on the underlying biological mechanisms of action (18, 23), it has informed effective dosing strategies for combination CAR T-cell and targeted radionuclide therapy (24), and CAR T-cell therapy in combination with the anti-inflammatory steroid Dexamethasone (25).

Despite the success of the CARRGO model, it is limited in the scope of potential phenomena that it can capture in regards to the precise interactions between the CAR T-cells and glioma cells. In this work, we use the SINDy modeling framework to incorporate important extensions to the CARRGO model. These extensions are: predator growth that is dependent on the density of prey, also known as a functional response (26, 27); individual predator and prey growth that saturates at some maximum value (logistic growth) (18), or has a population threshold below which collapse occurs (the Allee effect) (28, 29); and predator-prey interactions in which one or two CAR T-cells are bound to a single cancer cell at once, referred to as single or double binding, respectively (23, 30). Other efforts of extending CAR T-cell modeling have looked at fractional order derivatives (31) and stochastic dynamics (32) in the context of CAR T-cell treatment for viral infections, specifically coronaviruses. However, our treatment focuses on integer order derivatives and deterministic dynamics.

An ever-present challenge to quantitative biologists is fitting a proposed model to experimental data, also known as parameter estimation or model inference. On one hand, quantitative biologists seek models that capture as much biological realism and complexity as possible. On the other hand, increasing model complexity increases the computational challenge to accurately, confidently, and expediently determine model parameter values. This approach is further complicated if a researcher chooses to compare competing or complementary models (33, 34). An alternative approach, examined in this paper, is to leverage newly developed methods rooted in data science and machine learning which identify the strength of individual mathematical terms as candidates for an explanatory model. These methods are often referred to as dynamic mode decomposition, symbolic regression, or sparse identification.

Dynamic mode decomposition (DMD) is a data driven technique that interrogates time-series data by performing a singular value decomposition (SVD) on carefully structured matrices of the given data (35, 13). In this formalism, the orthonormal basis vectors generated by singular value decomposition serve as linear generators of the system dynamics such that forward prediction can be performed absent a known underlying mathematical model. Alternatively, SINDy identifies the specific mathematical terms that give rise to the observed dynamics governed by ordinary and partial differential equation models (15). SINDy achieves this by regressing experimental data onto a high-dimensional library of candidate model terms, and it has proven successful in climate modeling (36), fluid mechanics (37), and control theory (38). Since the initial publication of SINDy, several extensions have been studied, including: discovery of rational ordinary differential equations (39, 40); robust implementation with under-sampled data (41) or excessive noise (42); or incorporation of physics informed neural networks when particular symmetries are known to exist (43).

In its original and subsequent implementations, the CARRGO model demonstrated valuable utility in quantifying CAR T-cell killing dynamics when treating glioblastoma. Inferences of the underlying biological dynamics were made by examining how model parameter values changed along gradients of effector:target (E:T) ratios or as a function of other combination therapy concentrations. This is in direct contrast to the SINDy methodology, where the discovery of different model terms provides insight into the underlying biological dynamics as a result of variation along the E:T gradient. Here we compare of these two modelling frameworks on the same data set to provide further insight into the trade offs of *data first* versus *model first* approaches.

In this paper we utilize our experimental data to test these and other aspects of the DMD and SINDy frameworks. In Section 2.2 we introduce the families of models that are anticipated to be simultaneously biologically relevant and identifiable by SINDy, and we introduce a new approach to performing SINDy-based model inference. In Section 2.3.1 we present the latent variable analysis based on DMD that is used to generate the time-series CAR T-cell trajectories based on those of the cancer cells and the known boundary values for the CAR T-cells. In Section 2.3.2 we introduce the SINDy methodology in the particular context of our application. Results of our approach are presented in Section 3 where we (1) highlight how the discovered models vary as a result of different initial conditions in the cancer cell and CAR T-cell populations and (2) examine how well the discovered models found in this *data first* approach compare to a typical *model first* in characterizing the experimental data. In Section 4 we demonstrate how our results can guide experimental design to validate the predictions made by the discovered models, and we elaborate on some of the challenges encountered in this study.

## 2 MATERIALS AND METHODS

### 2.1 Experimental setup

The data analyzed in this study come from previously conducted experiments whose procedures are described in Sahoo et al. (18) and Brummer et al. (25), and summarized in Figure 1. The primary brain tumor cell line studied (PBT128) was selected for its endogenous high and relatively uniform expression of IL13R*α*2 antigen (89.11% IL13R*α*2+) (25). This cell line was derived from glioblastoma tumor resection tissue as described in (44, 45). To generate IL13R*α*2-targeted CAR T-cell lines, healthy donor CD62L+ naive and memory T-cells were lentivirally transduced to express second-generation 4-1BB-containing CAR that utilizes the IL13 cytokine with an E12Y engineered mutation as the IL13R*α*2 targeting domain (46).

**Figure 1.**
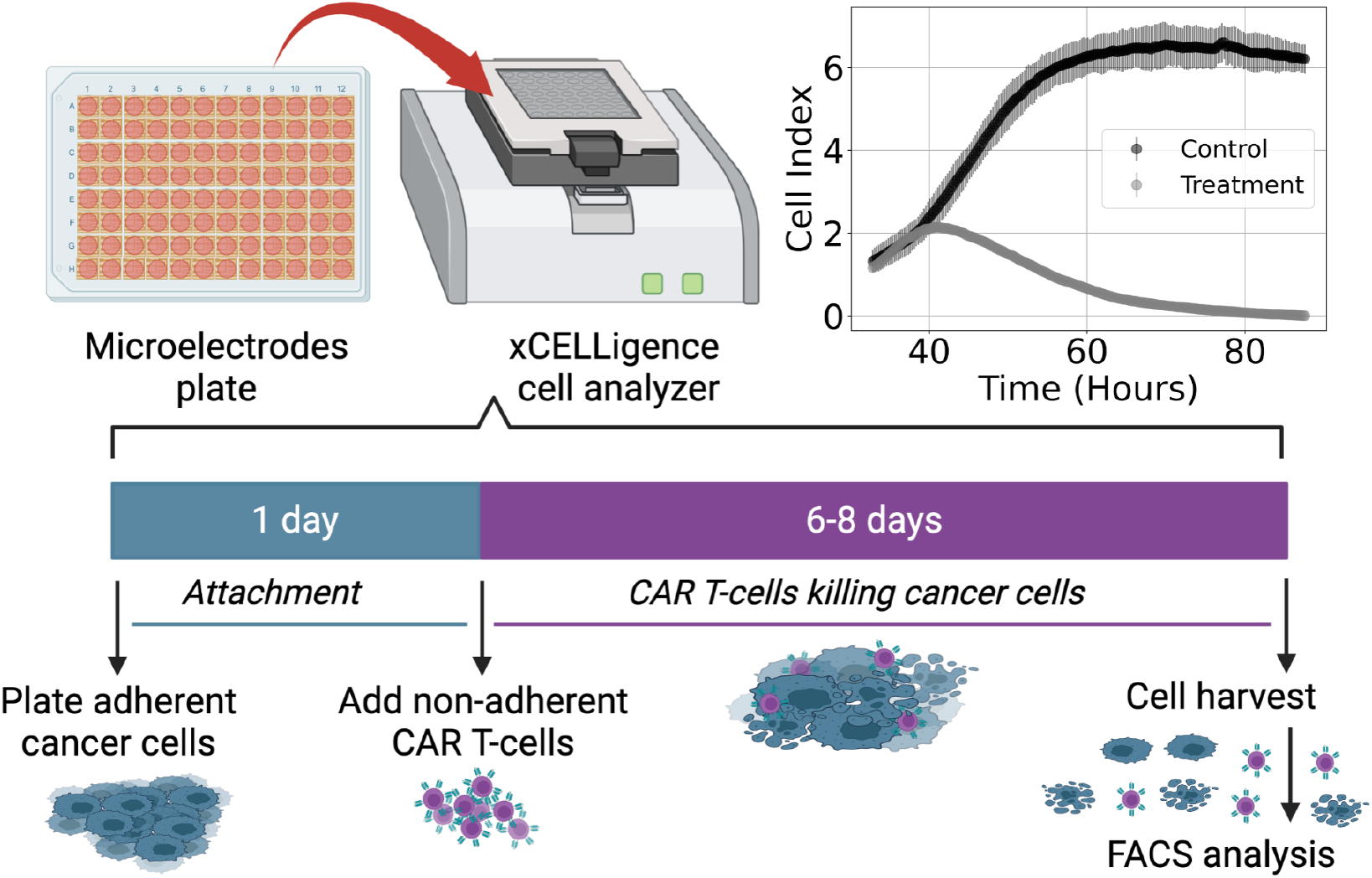
Diagram of experimental procedure highlighting use of microelectrode plates in an xCELLigence cell analyzer system and sample Cell Index (CI) measurements for control and treatment groups (E:T = 1:4). This system utilizes real-time voltage measurements to determine CI values representative of the adherent cancer cell population as a function of time. CAR T-cells are added following 24 hours of cancer cell expansion and attachment. After 6-8 days of monitoring the cancer cell growth and death dynamics, cells are harvested and enumerated using flow cytometry.

Cell killing experiments were conducted and monitored with an xCELLigence cell analyzer system. Measurements of cancer cell populations are reported every 15 minutes through changes in electrical impedance as cancer cells adhere to microelectrode plates, and are reported in units of Cell Index (CI), where 1 CI ≈ 10*K* cells (47, 48, 49). Flow cytometry was used to count the non-adherent CAR T-cells upon termination of the experiment. Measurements of CAR T-cell populations are reported in units of CI for the purposes of working in a common scale. We used the conversion factor of 1 CI ≈ 10*K* cells. Cancer cells were seeded at 10*K* − 20*K* cells and left either untreated or treated with only CAR T-cells, with treatments occurring 24 hours after seeding and monitored for 6-8 days (Figure 1). CAR T-cell treatments were performed with effector-to-target ratios (E:T) of 1:4, 1:8, and 1:20. All experimental conditions were conducted in duplicate.

### 2.2 Effective interaction models

Challenges to the *model first* approach to systems biology are (1) deciding on a sufficiently comprehensive model that captures all pertinent phenomena and (2) fitting the selected model to available data. Researchers are tasked with justifying their decisions in selecting candidate models. Yet, a common feature of dynamical systems models are the presence of ratios of polynomials. Such terms in ODEs can be difficult for the convergence of optimization algorithms to global solutions due to the possible existence of multiple local solutions within the model parameter space (50). In such instances researchers must either rely on high performance computational methods, have collected a vast amount of experimental data, or both. To address this problem, we utilize binomial expansions of candidate model terms under the assumptions of CAR T-cell treatment success and fast, irreversible reaction kinetics. In the following sections we present the space of possible models anticipated to characterize our experimental system, and the steps necessary to reduce the complexity of these candidate models.

The dynamical model that our experimental system is anticipated to follow is defined generically as,

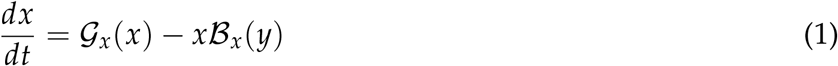

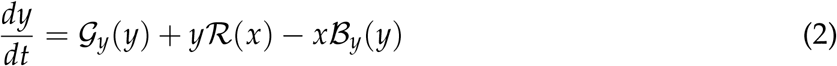

where 𝒢_*x*_ and 𝒢_*y*_ represent a growth-death model for the cancer cells, *x*, and the CAR T-cells, *y*. ℬ_*x*_ and ℬ_*y*_ represent a binding model for whether single or pairs of CAR T-cells attack individual cancer cells, and ℛ represents a model for the CAR T-cell functional response. In the subsections below, we explore different families of models representing the terms in the above equations. Explicitly, we examine different types of (a) Growth and death models, (b) Functional response models, and (c) CAR T-cell-cancer cell binding models.

#### 2.2.1. Growth and Death

We consider three different growth-death models for both the cancer cells and CAR T-cells. These are logistic growth, and the weak and strong Allee effect models, presented as,

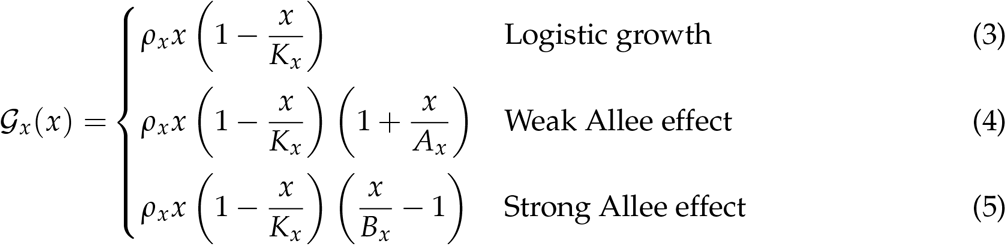

for 𝒢_*x*_(*x*), and similarly for 𝒢_*y*_(*y*). Here, *ρ*_*x*_ is the net growth rate, *K*_*x*_ is the population carrying capacity, *A*_*x*_ is a weak parameterization of deviations from logistic growth, and *B*_*x*_ is the threshold for population survival or death absent predation. All model parameters are assumed positive, with the added constraint that *K*_*x*_ *> B*_*x*_ *>* 0. We anticipate similar growth models for the CAR T-cells, 𝒢_*y*_(*y*), with allowance of different models for the different cell types and model constants. Logistic growth is commonly favored for its simplicity in experimental systems (18, 24, 25), while there is growing evidence that Allee effects are required for accurate characterization of low density cancer cell populations (28, 29, 51, 52) or as the result of directed movement (53), the latter of which being an observable feature of CAR T-cell behavior using bright field imaging (18, 25).

In Figure 2, graphs of population growth rates versus population size and population size versus time are presented for each growth model and for a variety of initial conditions. Parameter values used were *ρ* = 0.75 hrs^-1^, *K* = 10 CI, *A* = 5 CI, and *B* = 5 CI. Examination of the logistic growth model in Figure 2**a** and the weak Allee effect in Figure 2**b** demonstrates similar population saturation at the carrying capacity *K* = 10 CI, but a slight deviation between how the models reach saturation. Specifically, the weak Allee effect exhibits a reduced per capita growth rate at low population densities comapared to logistic growth. Examination of Figure 2**c** demonstrates the crucial difference between the strong Allee effect and either of the logistic growth or weak Allee effect through the existence of a minimum population threshold, *B*, above which the population will persist, and below which the population will die off.

**Figure 2.**
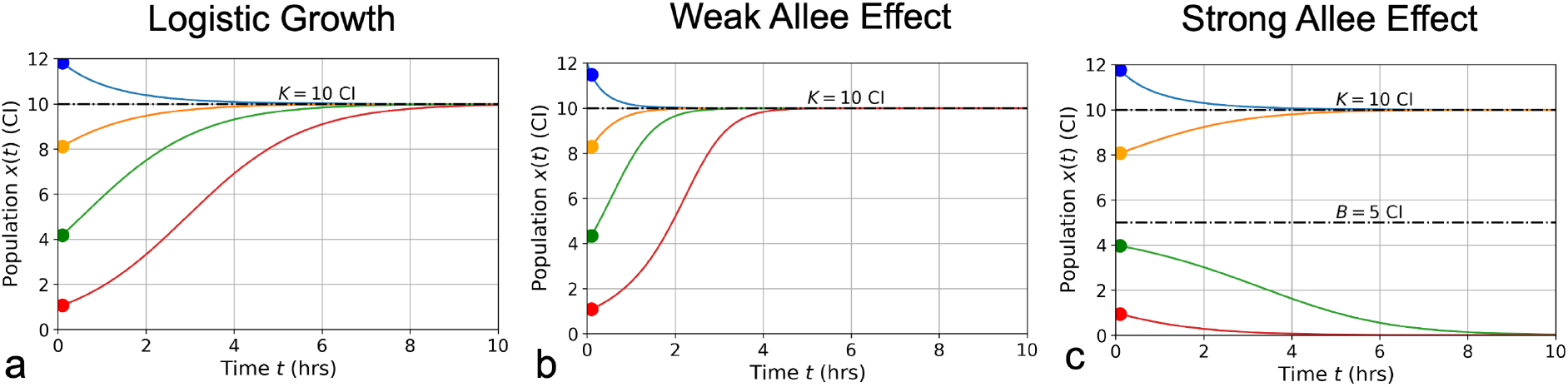
Conceptual graphs of population size (in cell index - CI) versus time (in hours - hrs) for the three growth models presented in Eqs. (3)-(5): logistic growth **a**; weak Allee effect **b**; strong Allee effect **c**. Model parameter values are: *ρ* = 0.75 hrs^-1^, *K* = 10 CI, *A* = 5 CI, and *B* = 5 CI. Colors correspond to different initial cancer cell seeding conditions which are the same for each model in a cancer cell only scenario (blue = 12CI, orange = 8CI, green = 4CI, red = 1CI).

Due to the fact that SINDy produces discovered models in their polynomial form without factoring, or grouping of terms together, we must consider the un-factored polynomial form of each model. To determine appropriate constraints on the model coefficients, we will expand the growth models and factor by common monomials. Doing so for 𝒢_*x*_(*x*) and dropping the subscript gives the following,

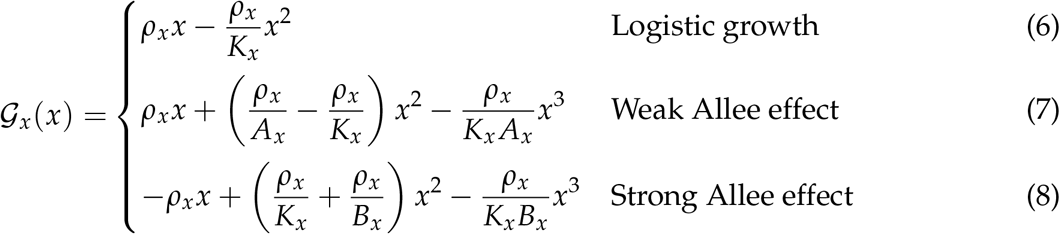

and similarly for 𝒢_*y*_(*y*). Here we can see that the coefficients for *x* and *x*^2^ can be positive or negative, but the coefficients for *x*^3^ must be fixed as negative values, where we have absorbed the minus signs in Eqs. (7)-(8) into *ρ*_*x*_/*K*_*x*_ *A*_*x*_.

#### 2.2.2 CAR T-cell-cancer cell binding

Cell binding models characterize the rates of formation and disassociation of conjugate pairs of species, also referred to as interaction molecules (Figure 3**a**). These models historically are known as Hill-Langmuir functions for their originating studies in hemoglobin formation (54) and gas adsorption on material surfaces (55), yet perhaps are better known for their use in modeling enzyme reaction kinetics, or Michaelis-Menten kinetics (56). The same modeling principles have been extended to examine cell binding in T-cell and cancer cell interactions (2, 23, 30). An important challenge to the field of cancer immunotherapy modeling is characterizing higher-order cell binding dynamics. That is, the formation of conjugates that consist of multiple CAR T-cells attacking single cancer cells (Figure 3**a**). These cancer cell-CAR T-cell conjugates are hypothesized to form as either a consequence of increased effector to target ratios or as a result of increased antigen density on target cells. As our experiment uses one single cell line with a high and uniform antigen expression level of IL13R*α*2, we assume on average all cancer cells have approximately the same antigen density. We thus focus our attention to experimental variation in the effector to target ratios.

**Figure 3.**
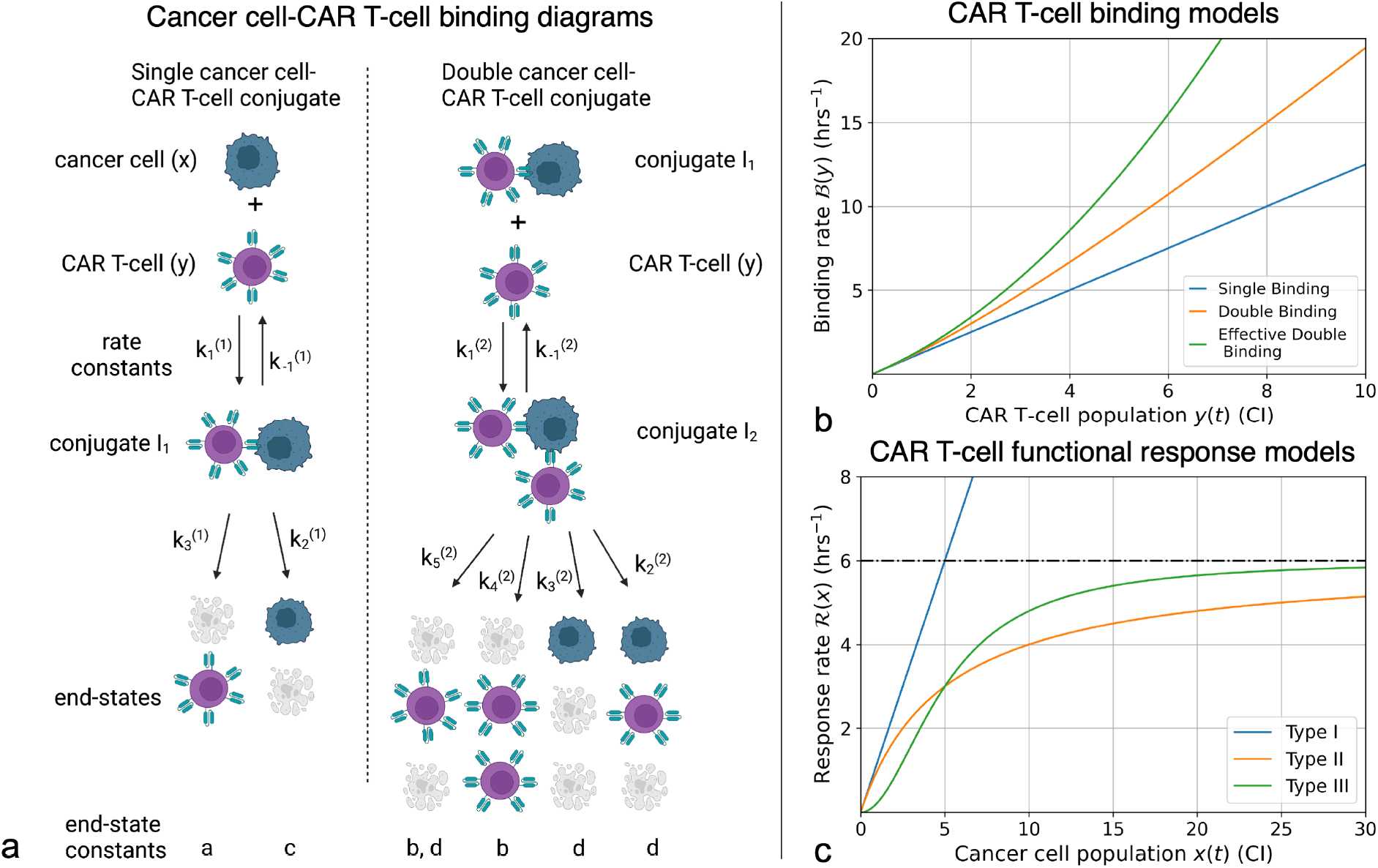
(**a**) Compartmental model for single and double CAR T-cell-cancer cell binding. Expressions for how rate constants 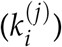 contribute to the growth or death of the cancer cell and CAR T-cell populations are presented in Eqs. (9)-(14). See (30) for further development and analysis of the cell binding model. (**b**) Graphs of binding rate versus CAR T-cell population for the single binding, double binding, and effective double binding models in Eqs. (9)-(12), (16), and (18). Model parameters for antigen bindings are: *a* = 20 CI^-2^·hrs^-2^ and *h* = 16 CI^-1^·hrs^-1^ for single binding; *a* = 20 CI^-2^·hrs^-2^, *b* = 5 CI^-3^·hrs^-2^, *h* = 16 CI^-1^·hrs^-1^, and *k* = 2 CI^-2^ hrs^-1^ for double binding; and *a* = 20 CI^-2^·hrs^-2^, *b* = 2.75 CI^-3^·hrs^-2^, *h* = 16 CI^-1^·hrs^-1^, and *k* = 2 CI^-2^·hrs^-1^ for effective double binding. These parameter values were chosen to highlight how well the effective double binding model can approximate both the single and double binding models at low CAR T-cell population values, *y <* 1 CI. Note that since the original double binding model in this scenario is concave-up, the effective double binding model parameters should be chosen to match concavity. This requirement sets a positivity constraint on the quadratic term in Eqs. (16) and (18). (**c**) Graphs of CAR T-cell response rates versus cancer cell population for different functional response models. Model parameters for functional responses are: *p* = 6/5 CI^-1^·hrs^-1^ for Type I; *p* = 6 CI^-1^·hrs^-1^ and *g* = 5 CI for Types II and III. Note overlap of Types I and II functional responses for *x <* 1 CI, and distinct differences in concavity between Types II (negative) and III (positive) for *x <* 2 CI. These characteristics correspond to Type I and Type II functional responses being indistinguishable at low cancer cell populations, and Type II and Type III being differentiated by fast-then-slow response rates (Type II) versus slow-then-fast response rates (Type III).

Following the work of Li et al. (30), we incorporate *fast irreversible* single and double cell binding into our generic model landscape. Here, fast binding implies that conjugate formation and dissociation occur quickly enough to maintain equilibrium in the conjugate populations, *I*_1_ and *I*_2_, such that *dI*_1_/*dt* = 0 and *dI*_2_/*dt* = 0. While irreversible means that all conjugate formation leads to death, or 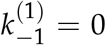 and 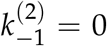. These assumptions are consistent with the conditions of relatively higher effector to target ratios, or high antigen densities on target cells. They also imply that a mixture of conjugates and dissociates may exist, but that the dynamics happen such that the conjugate populations are fixed and do not change with time. Furthermore, we only consider the higher-order binding scenario of two CAR T-cells to one cancer cell. Solving for the contributions to the cancer and CAR T-cell populations due to binding dynamics results in,

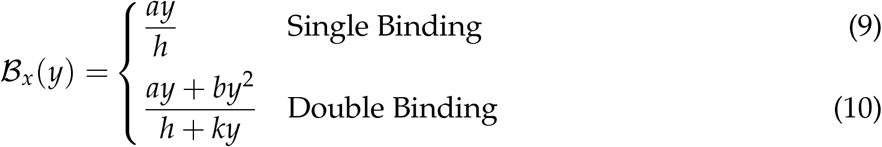

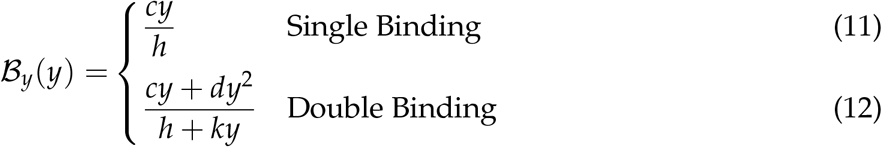

where the constants *a, b, c*, and *d* are defined in terms of the association rate constants, 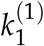 and 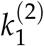, and the death rate constants 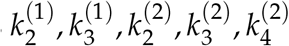 and 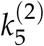 from Figure 3**a** as follows,

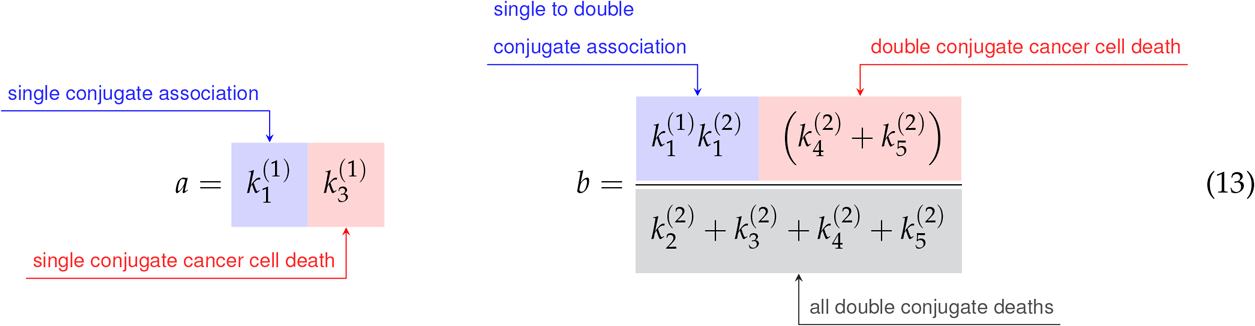

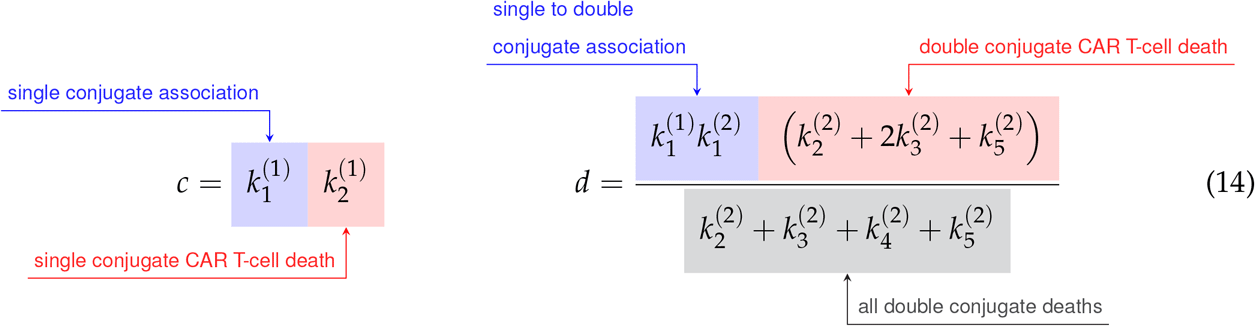

Finally, the constant *h* is the sum of the single conjugate death rates, 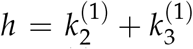, and the constant *k* is simply a renaming of the double conjugate association rate, 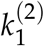. As the variable renaming is admittedly complicated, the constants *a, b, c*, and *d* are defined to quickly identify end states of conjugate formation and have been located next to their corresponding interaction products in Figure 3**a**.

The per-cancer cell binding models are graphed in Figure 3**b**. Model parameter values used for the single and double cell binding models in Eqs. (9)-(18)) are: *a* = 20 CI^-2^·hrs^-2^, and *h* = 16 CI^-1^·hrs^-1^ for single binding; and *a* = 20 CI^-2^·hrs^-2^, *b* = 5 CI^-3^·hrs^-2^, *h* = 16 CI^-1^·hrs^-1^, and *k* = 2 CI^-2^·hrs^-1^ for double binding. We highlight that we are restricting ourselves to scenarios where increases in the CAR T-cell population during a given trial leads to increases in the likelihood of double binding, which results in super-linear increase of per-cancer cell binding. This restriction enforces concavity of the effective double cell binding model which we explore next. It is possible for the double binding model to exhibit a sub-linear increase in per-cancer cell antigen binding as the CAR T-cell population increases, and an overall decrease in cancer cell killing. However, this scenario does not agree with our experimental data of increased killing with increased effector-to-target ratios.

Importantly, the rational forms of the binding rates typically complicate determination of parameter values in conventional dynamical modeling. To reduce model complexity, we take advantage of potential differences between the rates of conjugate association and conjugate death that can give rise to simplifications. If the product of the CAR T-cell population and the rate of forming double conjugates, *ky*, is small compared to the sum of the rates of single conjugate deaths, *h*, then *ky*/*h <* 1, and we can again perform a binomial expansion in the cell binding denominators. A second way of interpreting this condition is to require the number of CAR T-cells to remain small compared to the ratio of the rate of double conjugate formation to the sum of the rates of single conjugate deaths, *y < h*/*k*. Performing the binomial expansion and truncating again at 𝒪(*y*^2^) results in the following effective models of cell binding,

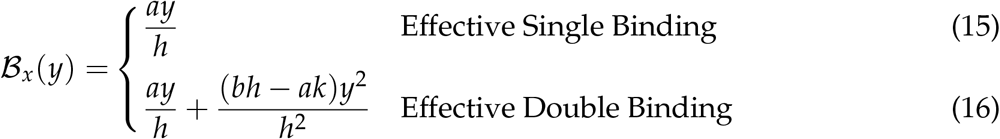

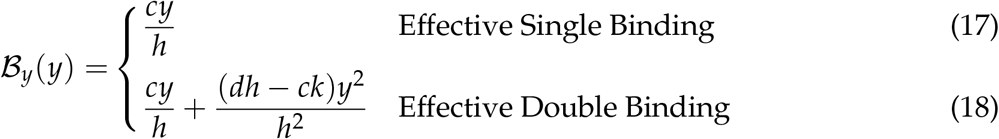

Here the effective double conjugate antigen binding model takes the form of the exact single conjugate binding model plus a correction due to double conjugate formation. Eqs. (16) and (18) are graphed in Figure 3**b**, using the parameter values of *a* = 20 CI^-2^·hrs^-2^, *b* = 2.75 CI^-3^·hrs^-2^, *h* = 16 CI^-1^·hrs^-1^, and *k* = 2 CI^-2^·hrs^-1^. These values are chosen to demonstrate that the effective double binding model can accurately approximate both the exact single and double binding models for small CAR T-cell populations, *y <* 1 CI. Importantly, we note that if the parameter values *b* or *d* are sufficiently small, corresponding to low double conjugate CAR T-cell or cancer cell death rates, then the quadratic terms in Eqs. (16) and (18) will be negative, and the concavity of the effective double binding model deviate significantly from the exact model. This phenomenological consideration of the effective models sets an important constraint on the positivity of the coefficients for the quadratic terms in Eqs. (16) and (18), which we will revisit in Section 2.3.

#### 2.2.3 Functional response

We next consider the first three types of functional response models that characterize how the CAR T-cells respond, or expand, in the presence of cancer cells. These models are defined as,

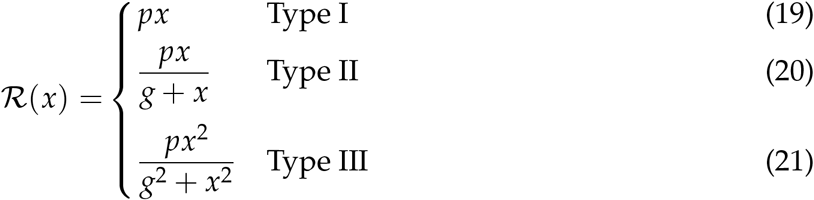

where *p* is the predator response, or CAR T-cell response rate, and *g* is the prey population density threshold at which predator behavior changes (e.g. fast-to-slow or slow-to-fast rates of killing). Functional responses model changes in predator hunting due to the prey density, generally defined with respect to some prey population threshold, here denoted as *g*. The population dependence on predator hunting behavior can also be interpreted as a handling time for distinguishing between time spent seeking prey, or recognizing cancer cells, and time spent consuming and attacking prey (26, 27, 19).

The three types of functional responses are graphed in Figure 3**c**. In a Type I functional response, the predator response is constant for all prey population sizes. The interpretation of this response is that there are no differences in time or cost between all predator functions (searching and capture). In a Type II functional response the predator response is linear at low prey density (mirroring a Type I behavior) yet saturates at high prey density. Finally, in a Type III functional response the predator response is low at low prey densities, reflecting the potential for cancer cells to escape immune surveillance, yet again saturates at high prey densities, with a linear response at intermediate prey densities.

As with the binding rate models, the rational forms of Types II and III functional responses present challenges to model discovery methods. Thus, we assume a significant level of effectiveness in CAR T-cell treatment such that the cancer cell population remains relatively low with respect to the functional response threshold, that is *x < g*, or *x*/*g <* 1. CAR T-cell effectiveness is demonstrated in Figure 1, where the control cancer cell population is shown to achieve a maximum population of approximately 6.5 CI, while the treatment population of *E* : *T* = 1 : 4 reaches a maximum population of approximately 2CI. The approximation condition permits the use of a binomial expansion about *x* = 0 on the denominators for the Types II and III functional responses, resulting in,

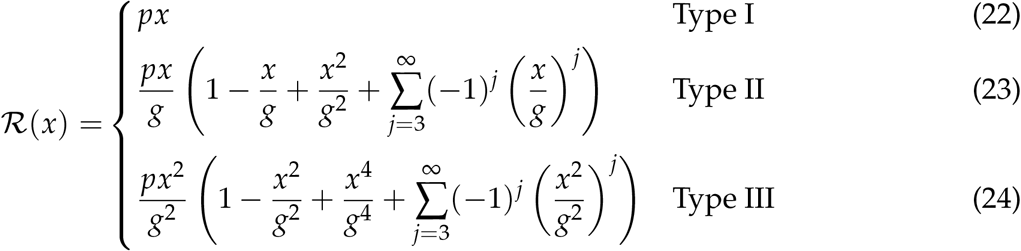

Further assuming that contributions to the functional response models of 𝒪(*x*^3^/*g*^3^) or greater are negligible, we terminate the expansions at 𝒪(*x*^2^/*g*^2^) to arrive at the following effective functional response models,

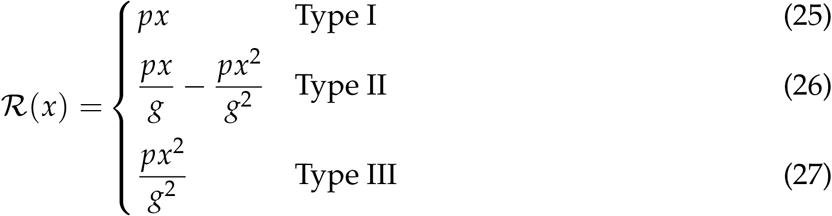

It is important to highlight that the leading order term for the expansion for a Type II functional response is indistinguishable from a Type I functional response. This feature is reflected by the overlap in the graphs of the Type I and Type II responses presented in Figure 3, where the cancer cell population is small, *x* ∈ [0, 1] CI, compared to the value of *g* = 5 CI. As the cancer cell population increases, the density dependence of the CAR T-cells starts to take effect as demonstrated by the parabolic contribution of the Type II response. In contrast to this, the expansions for functional responses of Types II and III are significantly unique from one another. Specifically, only expansions for Type II can lead to odd-powered terms in *x*, and although both expansions can express similar even-powered terms, they come with different concavities. That is, at small cancer populations the Type II functional response is characterized as a concave down parabola, while the Type III functional response is characterized as a concave up parabola. This difference regarding the positivity of the terms that are of second-order dependence in *x* corresponds to the different density dependent behaviors of the CAR T-cells at small cancer cell populations, specifically that Type II is a fast-to-slow response rate while Type III is a slow-to-fast response rate.

By performing the approximations used to derive Eqs.(26)-(27), and using truncated terms, we have reduced the complexity of the functional response terms. This step will simplify the process of model discovery. However, since this step assumes that the prey population remains small compared to the functional response threshold, the number of terms needed in Eqs.(23)-(24) for accurate characterization of the system dynamics may vary as a result of experimental variation in the effector to target ratio of the CAR T-cells and the cancer cells. This variation in the effector to target ratio may also influence the structure of other interaction terms, specifically those pertaining to the single or paired binding dynamics.

#### 2.2.4 Landscape of effective models

To gain a broader perspective of the overall form of our ODE models, we substitute the effective models for functional responses and antigen binding into Eqs. (1)-(2), arriving at,

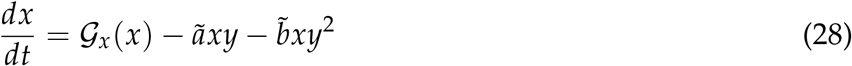

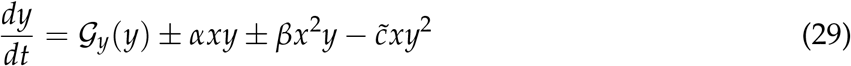

where 𝒢 again represents any of the potential growth-death models under consideration, *ã* = *a*/*h* and 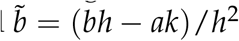 are redefined constants (both assumed to be positive) for the coefficients of the effective single and double binding models for the cancer cells, *αxy* = (*p*/*g* − *c*/*h*)*xy* and represents the combination of first order terms for CAR T-cell response and single binding, *βx*^2^*y* = (*p*/*g*^2^)*xy* and represents the potential second order term from the CAR T-cell response, and 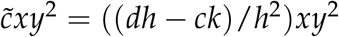 represents the effective double binding model for the CAR T-cells. We have explicitly used ± notation to indicate that we do not know *a priori* the signs for the *xy* and *x*^2^*y* terms in Eq. (29), as these are determined by the relative contributions of Type I and first order Type II-like CAR T-cell responses and single antigen binding for the *xy* term, and whether or not second order Type II or first order Type III CAR T-cell response is occurring for the *x*^2^*y* term. The benefit of the approach demonstrates the presence and/or sign conventions of the various model coefficients that we determine using the SINDy model discovery algorithm can be directly interpreted in terms of different underlying biological phenomena.

### 2.3 Model discovery

Our implementation of the model discovery techniques of dynamic mode decomposition and sparse identification of non-linear dynamics (SINDy) is performed in two stages. First is latent variable analysis, the extraction of the latent variable representing the CAR T-cell population from the time-varying cancer cell population. The second step is implementation of SINDy, whereupon the functional terms of the underlying models describing the dynamical system are determined.

#### 2.3.1 Latent Variable Analysis

Despite having only measured the initial and final CAR T-cell populations, we can utilize latent variable analysis to infer the hidden CAR T-cell dynamics from the cancer cell dynamics. We do this using the delay coordinate embedding of Taken’s theorem to reconstruct the attractor of the system that is known to exist in more dimensions than those measured (13, 15, 57). The first step in this approach is to assemble a Hankel matrix, **H**, by stacking delayed time-series of the cancer cell measurements *x*(*t*) as follows,

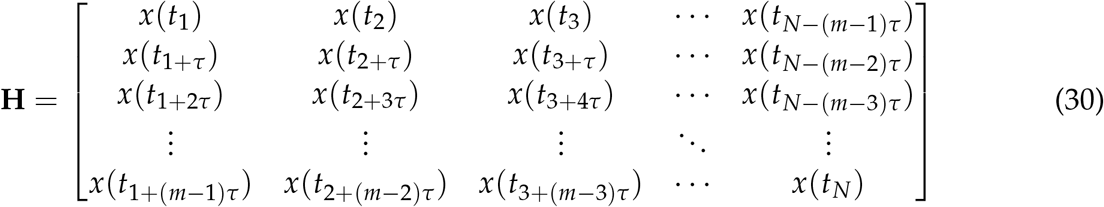

where *τ*, known as the embedding delay, represents the size of the time-delay we use, and *m*, known as the embedding dimension, represents both the number of rows that we assemble in the Hankel matrix and, importantly, the number of anticipated latent variables we expect to find.

To minimize the effects of experimental noise on the results of Taken’s Theorem, we splined our cancer cell trajectories and re-sampled at the same experimental sampling rate of one measurement per 15 minutes. The function *smooth*.*spline* from the programming language **R** was used to perform the splining. This function uses cubic splines to approximate trajectories, with a penalty term to control for trajectory curvature. The number of knots used to spline each trajectory were determined by inspection, and are recorded in the analysis code available at https://github.com/alexbbrummer/CART_SINDy. Further details on the splining methods used are available in (58).

To determine optimal values for *τ* and *m*, we can use two separate formulae to inform the decisions (57). The optimal time delay is determined by the value of *τ* which minimizes the mutual information between measurements. This is done by dividing the interval [*x*_*min*_, *x*_*max*_] into *j* equally sized partitions, and calculating the probability *P*_*k*_ that a measurement of the time series is in the *k*^th^ partition, and the probability *P*_*h,k*_ that a measurement *x*_*i*_ is in the *h*^th^ partition while the neighboring measurement *x*_*i*+*τ*_ is in the *k*^th^ partition. Mutual information is given by

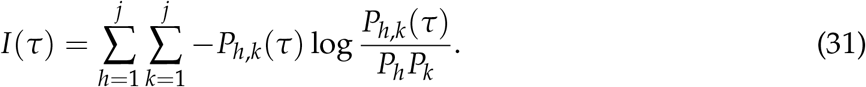

The optimal time-delay to use for a given time series is selected by finding the value of *τ* which results in the first minimum value in mutual information, or arg min|_*τ*_{*I*(*τ*)}. A graph of mutual information versus time delay is presented in Supplemental Figure **S1a**. For our cancer cell time series data, this optimal time delay value was found to be *τ* = 1.

To determine the embedding dimension, *m*, we calculate the number of false nearest neighbors to a given measurement as the time series is embedded in successively greater dimensional spaces. This calculation is done to ensure that the attractor constructed from the latent variables remains smooth upon embedding. We perform the calculation iteratively by starting with a point *p*(*i*) in an *m*-dimensional embedding, and identifying a neighboring point *p*(*j*) such that the distance between *p*(*i*) and *p*(*j*) is less than a constant value typically chosen as the standard deviation of the data. Next, the normalized distance between the points *p*(*i*) and *p*(*j*) in the *m* + 1-dimensional embedding is calculated using the following expression,

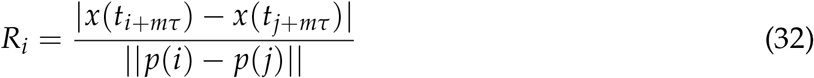

*R*_*i*_ is calculated across the entire time series and iteratively for greater embeddings, *m* = 1, 2, 3, …. False nearest neighbors are identified when *R*_*i*_ *> R*_*threshold*_, where *R*_*threshhold*_ = 10 has been identified as satisfactory for most datasets (57). The ideal embedding dimension *m* is finally determined as that which results in a negligible fraction of false nearest neighbors. In Supplementary Figure **S1b** we present the calculated fraction of false nearest neighbors versus embedding the dimension. For our dataset, we identified *m* = 2 as the ideal embedding dimension, indicating the existence of one latent variable that we interpret as representing the CAR T-cell population.

Using values of *τ* = 1 for the time delay and *m* = 2 for the embedding dimension results in the following form of the Hankel matrix,

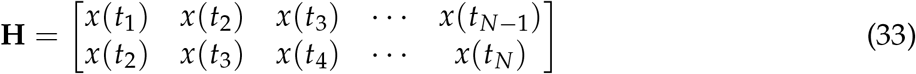

To extract the latent variable that represents the CAR T-cell time series, we perform a singular value decomposition of the Hankel matrix, **H** = **U**Σ**V**^∗^ (15, 13). Here, the columns of **V** represent scaled and standardized versions of both the original data in the first column, and approximations of the latent data in the subsequent columns. As our experimental procedure measured the initial and final CAR T-cell populations, our final step was to re-scale and offset the latent CAR T-cell variable extracted from the second column of **V**. We note that latent variable analysis is conducted on each trial for each experimental condition separately. In Figure 4 we present the measured cancer cells and CAR T-cells in addition to the discovered latent CAR T-cell time series for each effector to target ratio considered for the first of the two duplicate trials. In the Supplemental Material Figure **S2** we present the results of the latent variable analysis for the second of the two duplicate trials.

**Figure 4.**
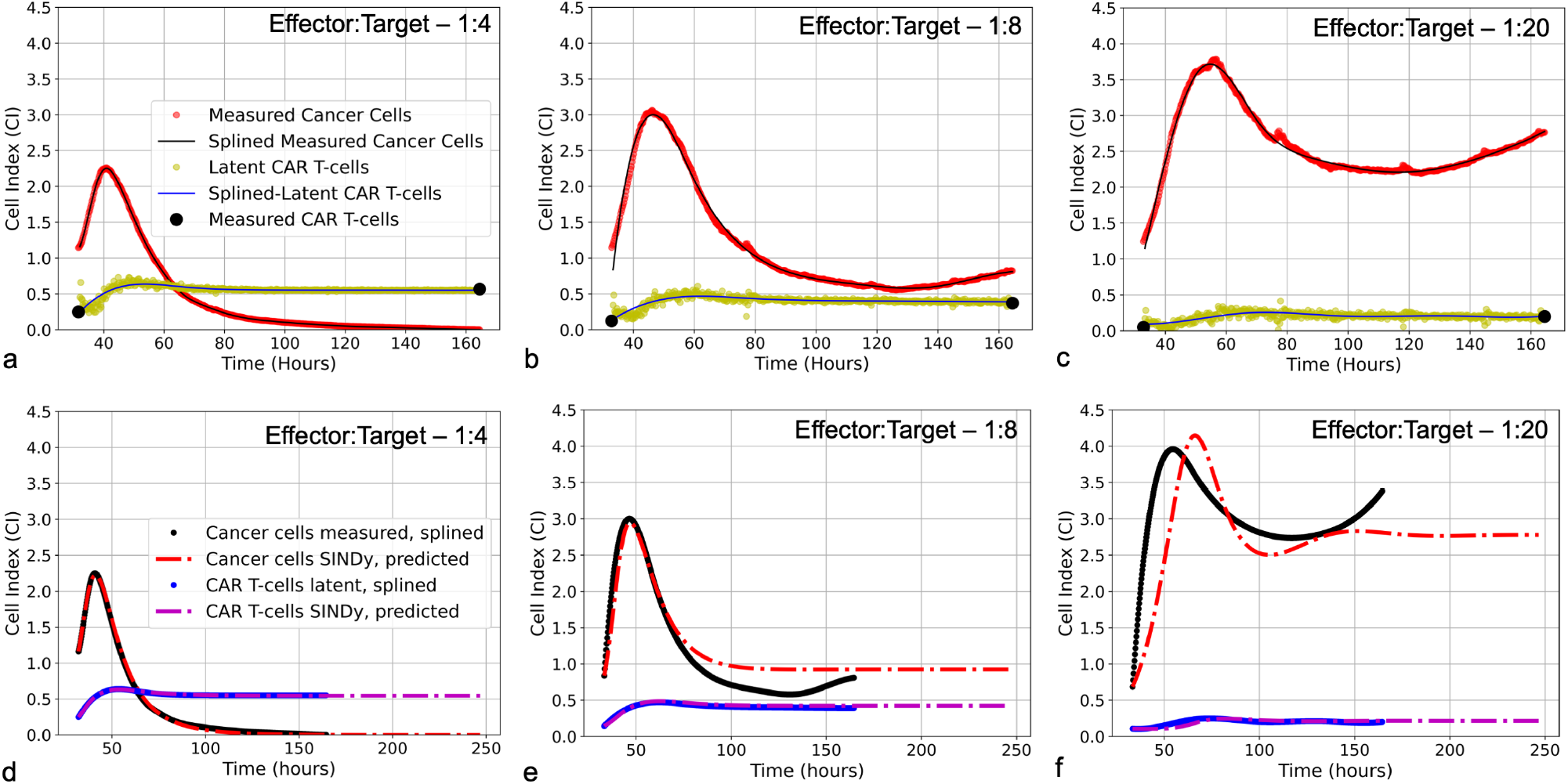
(**a-c**) Latent variable analysis results for first of two experimental replicates for each E:T ratio examined. Presented for are the cancer cell index measurements from the xCELLigence machine in red, overlaid with the splined measurements for the cancer cells in black; the two endpoint measurements for the CAR T-cell levels enumerated by flow cytometry in black, with the CAR T-cell population trajectory as determined by latent variable analysis in yellow, overlaid with the splined CAR T-cell trajectory in blue. Note that despite the CAR T-cell populations being measured with flow cytometry, we have converted levels to units of Cell Index for ease of comparison with the cancer cells, using a conversion factor of 1CI ≈ 10, 000 cells. (**d-f**) Predicted trajectories of discovered models compared to splined measurements of cancer cells and CAR T-cells for same data presented in (**a-c**). Splined cancer cell and CAR T-cell measurements are in black and blue, respectively. Predicted trajectories for cancer cells are the red dot-dashed lines, while the CAR T-cells are the purple dot-dashed lines. To examine stability of SINDy-discovered models, both simulations and forward predictions are presented to show steady-state behavior. Note that the best fits between predictions and measurements occur in the high E:T scenario, where assumptions made regarding treatment success and low cancer cell populations in determining model candidate terms are best adhered. As the E:T ratios get smaller, increasing deviation between discovered model predictions and splined measurements can be qualitatively observed. This is likely due to weakening of assumptions of treatment success and low cancer cell populations associated with the low E:T conditions. See Supplemental Material Figure **S2** for equivalent latent variable analysis results and SINDy-predicted trajectories for the second set of experimental replicates.

#### 2.3.2 Sparse identification of non-linear dynamics

SINDy is a data-driven methodology that discovers dynamical systems models through symbolic regression (15, 13). From a conceptual perspective, SINDy allows for the transformation of an analytical, first-order, non-linear dynamical systems model, expressed as

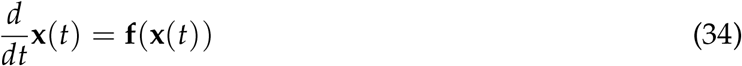

to a linearized matrix-model, expressed as

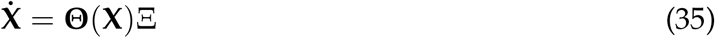

where 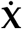 are numerical time-derivatives of our measured data, Θ(**X**) is a library of candidate functions that may describe the data and is evaluated on the measured data, and Ξ consists of the coefficients for the model terms from Θ(**X**) that describe the time-varying data 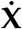. The objective of SINDy is to identify the sparsest version of Ξ, where sparsity is defined as the compromise between fewest number of non-zero terms with the greatest level of accuracy. In the context of our measurements for populations of cancer cells, *x*(*t*), and CAR T-cells, *y*(*t*), and the anticipated models for cell growth and interactions, 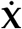 takes the following form,

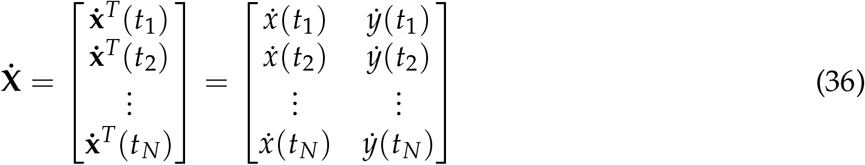

and **Θ**(**X**) is expressed as,

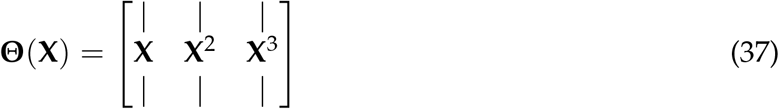

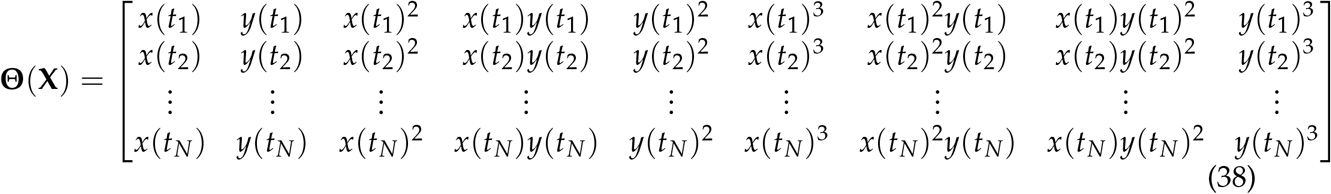

By solving the matrix-inverse problem in Eq. (35), we can find the column vectors Ξ that determine the coefficients for the model terms *ξ* that form the non-linear dynamical system best describing the measured data. To construct 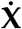 and **Θ**(**X**) from our duplicate trial experiments, the data from the repeated trials is stacked row-wise. Thus, only a single model will be discovered to explain all data for a given set of experimental conditions (e.g. effector-to-target ratios). Having repeat measurements is an important aspect for SINDy to converge on an accurate model, thus performing SINDy on averages of experimental trials undermines performance. For experimental conditions that have an abundance of experimental replicates, an AI-inspired division of data into training and testing sets can be conducted (59).

Once 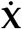 and **Θ**(**X**) have been constructed, a simple least-squares algorithm for solving Eq. (35) will result in a dense coefficient vector Ξ, thus we enforce sparsity of the coefficient vector Ξ through the method of sparse relaxed regularized regression (SR3) (60), where we seek optimization of the expression,

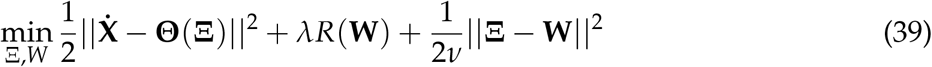

where **W** is the relaxed coefficient matrix that approximates **Ξ**, *R*(**W**) is the regularization of **W**, and *ν* and *λ* are hyper parameters that control how precisely **W** approximates **Ξ** and the strength of the regularization, respectively. For our problem, we chose to regularize under the *ℓ*1-norm with *ν* = 1 × 10^−5^. To determine the value of *λ*, we followed the approach taken in (39) in which we repeat the analysis for a range of *λ* values from *λ* ∈ [10^−8^, 10^1^] to calculate Pareto fronts between the root-mean-squared error between the measured and subsequently predicted values of **X** and the number of active terms from our library. In Supplementary Figure **S3** we present Pareto fronts for each of the experimental conditions for the varying effector to target ratios.

As discussed in Section 2.2, there are a variety of constraints we can expect for possible coefficients based on expected signs, or the absence of particular terms. An extension to SINDy allows for the incorporation of these constraints to ensure spurious terms are not discovered (61).

To make clear the constraints that were imposed, we can re-write Eq. (35) symbolically and in terms of the coefficients *ξ*_*i,j*_ as,

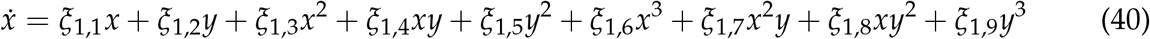

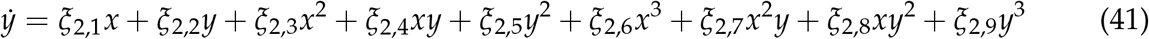

Then, the constraints that are imposed as per the anticipated effective models from Section 2.2 are,

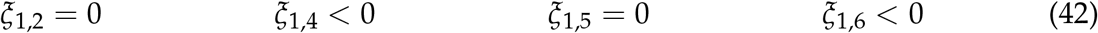

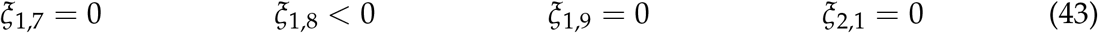

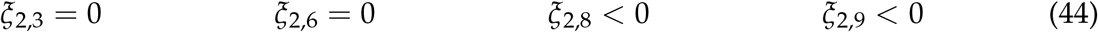

while the other 6 coefficients in *ξ*_*i,j*_ are left to freely vary.

Implementation of SINDy SR3 with constraints was performed using PySindy, a package designed for a wide array of implementations of the SINDy algorithm for spatio-temporal model discovery written in the programming language Python (16, 17). Included in the Supplemental Material are the associated datasets and Jupyter notebooks used for this study.

Finally, we highlight that the implementation of SINDy which we are relying on is designed specifically for explicit ordinary differential equations. An extension of SINDy exists for discovering ODEs with ratios of polynomials (39, 40), however this variation requires a significantly greater volume of data than that which we could collect. This is the underlying motivation behind our efforts to derive the effective models, thereby converting them into explicit ODEs and making effective usage of the volume of experimental data available by the study methods most usable for model discovery.

## 3 RESULTS

### 3.1 Discovered models and simulated comparison

Upon implementing SINDy on the CAR T-cell cancer cell killing data and performing the Pareto front analysis described in Section 2.3, we identified three distinct models describing the experimental data. Model selection is presented in Supplementary Figure **S3**, where we present the tradeoffs between model complexity, represented by the number of activated library terms, and either threshold *λ* or the root-mean-squared-error between the measured data and simulated data for each identified model. Our examination of the Pareto fronts found models with eight terms for E:T of 1:4, and 1:8, and a six term model for an E:T of 1:20. Below we summarize each of these models and in relation to how well they predict the measured data in Figure 4. We synthesize the coefficients and associated model categories for growth in Table 1 and for the CAR T-cell functional response and cell binding in Table 2.

**Table 1.**
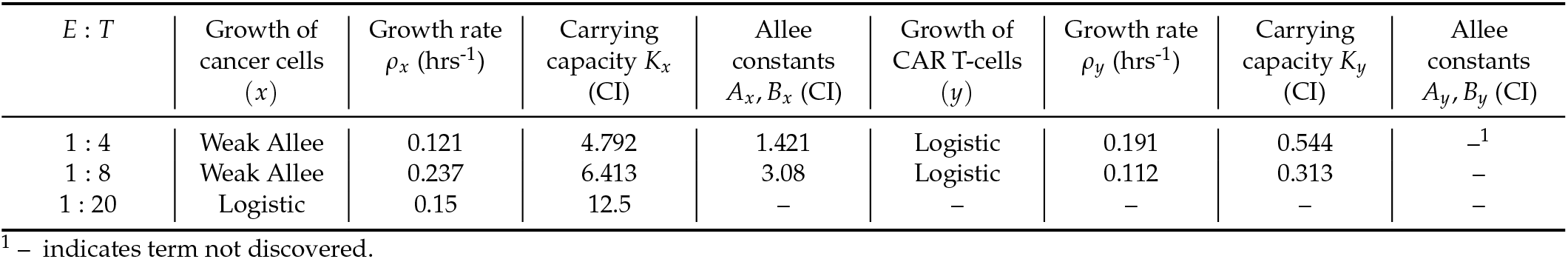
Coefficients for discovered growth model terms across all effector to target ratios.

**Table 2.** Coefficients for discovered interaction model terms across all effector to target ratios.

| $E : T$ | Response of<br>CAR T-cells | Type I & II<br>response $\alpha$<br>(CI<sup>-1</sup> hrs<sup>-1</sup>) | Type II & III<br>response $\beta$<br>(CI<sup>-2</sup> hrs<sup>-1</sup>) | Cancer<br>cell-CAR<br>T-cell binding | Single binding<br>$\tilde{a}$ (CI<sup>-1</sup> hrs<sup>-1</sup>) | Double<br>binding $\tilde{b}$<br>(CI<sup>-2</sup> hrs<sup>-1</sup>) | Double<br>binding $\tilde{c}$ (CI<sup>-2</sup><br>hrs<sup>-1</sup>) |
| --- | --- | --- | --- | --- | --- | --- | --- |
| 1 : 4 | Type II | 0.035 | -0.009 | Double | -<sup>1</sup> | 0.593 | - |
| 1 : 8 | Type II | 0.051 | -0.009 | Single | 0.626 | - | - |
| 1 : 20 | Type III | -0.002 | 0.005 | Mixed | 0.545 | - | 0.063 |

<sup>1</sup> - indicates term not discovered.

#### 3.1.1 High E:T discovered model

For the E:T = 1:4 data, the SINDy-discovered model takes the following form,

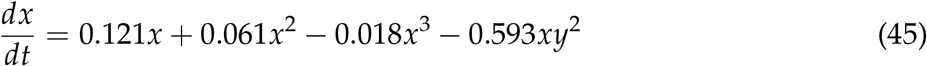

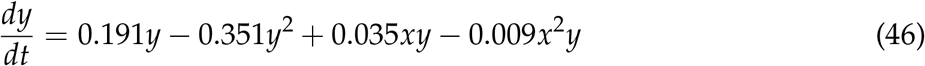

Factoring the terms related to single-species growth, we arrive at,

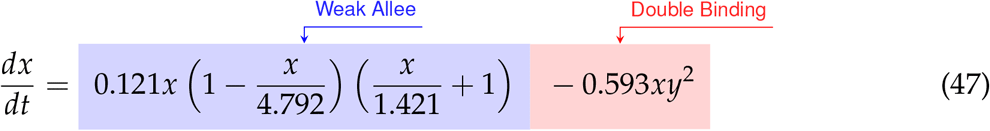

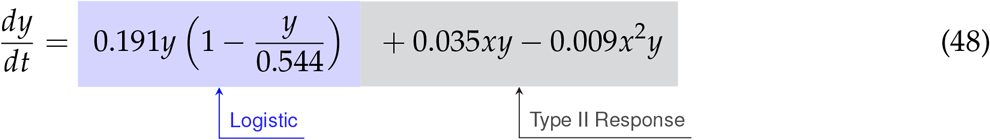

From Eqs. (47)-(48) we can interpret the discovered types of growth models and interactions. For cancer cell growth in Eq. (47), the observable structure indicates a weak Allee effect, with a growth rate of *ρ* = 0.121 hrs^-1^, a carrying capacity of *K* = 4.792 CI, and an Allee constant of *A* = 1.421 CI. For the CAR T-cells we find a logistic growth model with growth rate *ρ* = 0.191 hrs^-1^ and carrying capacity *K* = 0.544 CI. From the coefficients of *α* = 0.051 CI^-1^ hrs^-1^ on *xy* and *β* = −0.009 CI^-2^ hrs^-1^ on *x*^2^*y* for the CAR T-cells, we can infer a Type II functional response as the signs are positive and negative, respectively. Finally, the presence of an *xy*^2^ term in the cancer cells with a coefficient of 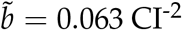 hrs^−1^ indicates the occurrence of double binding, notably in the absence of both the *xy* term in the cancer cells and the *xy*^2^ term in the CAR T-cells.

#### 3.1.2 Medium E:T discovered model

The SINDy-discovered model for the E:T = 1:8 data takes the following form,

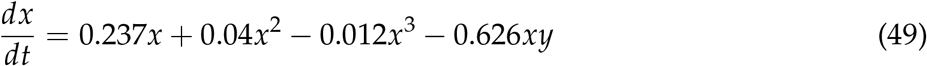

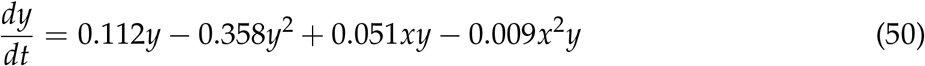

Factoring the terms related to single-species growth, we arrive at,

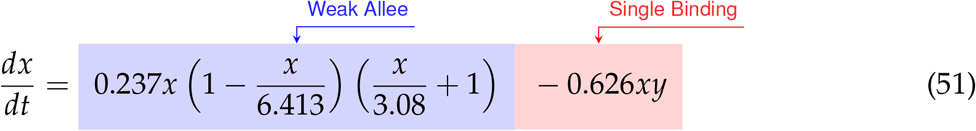

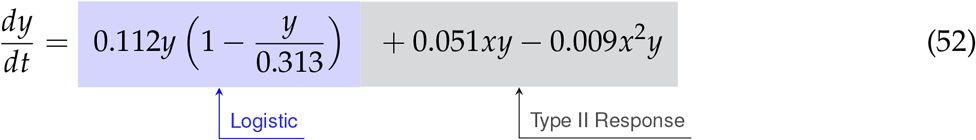

The model discovered for medium E:T is largely similar to that at high E:T. A weak Allee effect in growth is observed for the cancer cells, with growth rate *ρ* = 0.237 hrs^-1^, carrying capacity *K* = 6.413 CI, and Allee constant *A* = 3.08, while a logistic growth is observed for the CAR T-cells with growth rate *ρ* = 0.112 hrs^-1^ and carrying capacity *K* = 0.313 CI. We also observe a Type II CAR T-cell functional response, again indicated from the sign of the coefficients of *α* = 0.051 CI^-1^ hrs^-1^ and *β* = −0.01 CI^-2^ hrs^-1^ on the *xy* and *xy*^2^ terms being positive and negative, respectively. Unlike the high E:T scenario however, here we find evidence only of single binding from the sole presence of an *xy* term in the cancer cells with a coefficient of *ã* = −0.626 CI^-1^ hrs^-1^.

#### 3.1.3 Low E:T discovered model

Finally, for the E:T = 1:20 data the discovered model is,

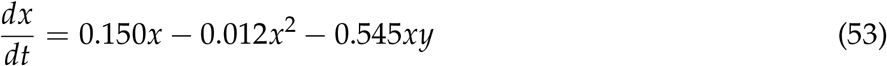

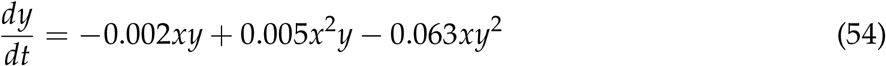

Factoring the terms related to single-species growth, we arrive at,

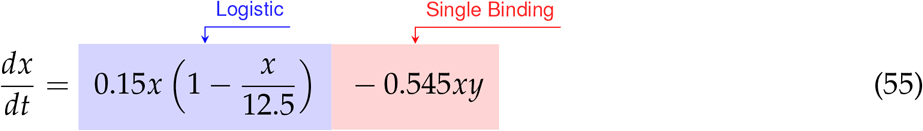

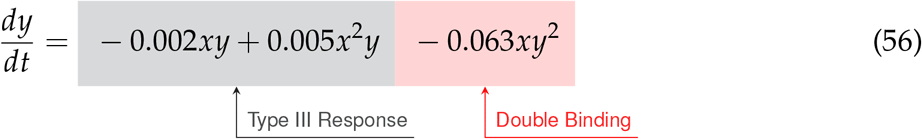

In this scenario we find significantly different growth and interaction models. The cancer cells show logistic growth, with growth rate *ρ* = 0.15 hrs^-1^ and carrying capacity *K* = 12.5 CI, while the CAR T-cells have no growth model. This time, as the signs for the coefficients of *α* = −0.002 CI^-1^ hrs^-1^ and *β* = 0.005 CI^-2^ hrs^-1^ on the *xy* and *x*^2^*y* terms for the CAR T-cells are now negative and positive, respectively, we infer a Type III functional response. Interestingly, we find a mixture of indicators for both single binding and double binding. This comes from the presence of only the *xy* term in the cancer cell model with a coefficient of *ã* = −0.545 CI^-1^ hrs^-1^, and of an *xy*^2^ term in the CAR T-cell model with a coefficient of 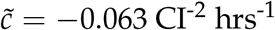.

All three E:T ratios of 1:4, 1:8, and 1:20 resulted in discovered models that accurately characterized the data, with root-mean-squared-errors of 0.02, 0.195, and 0.359, respectively. We highlight the discovery of consistent growth models of a weak Allee effect for the cancer cells and logistic growth for the CAR T-cells for the E:T ratios of 1:4 and 1:8. Importantly, the growth rates and carrying capacity for these scenarios were found to be comparable across E:T ratios. Interestingly, we observe a Type II functional response in the CAR T-cells functional response for both E:T = 1:4 and 1:8, and a transition to Type III for E:T = 1:20. Similarly, our discovered models indicate a transition from double to single binding as the E:T ratio changed from 1:4 to 1:8, and a model with mixed single and double binding terms was discovered for the E:T = 1:20.

### 3.2 Comparison with CARRGO model

We compared the *data first* model discovery methodology of SINDy against the CARRGO model, a traditional *model first* approach originally used to analyze and interpret the CAR T-cell killing dynamics (18, 25). The CARRGO model is defined as,

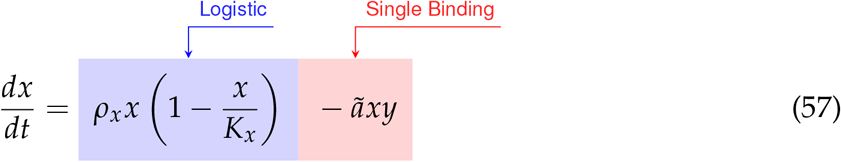

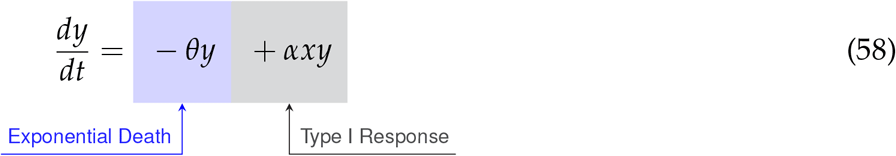

where we have expressed the parameter variables of the CARRGO model in terms of those used in the SINDy model for ease of comparison. From here we can see that the CARRGO model assumes logistic growth in the cancer cells, single binding between the cancer cells and CAR T-cells, a Type I functional response in the CAR T-cells, and exponential CAR T-cell death.

In Figure 5 are graphs of the best-fit versions of both the CARRGO model and SINDy discovered models for each E:T ratio. These fits were performed using the Levenberg-Marquadt optimization (LMO) algorithm, which requires initial guesses and bounds for each model parameter value. For the CARRGO model published parameter values were used for the starting guesses, while for the SINDy discovered models the discovered parameter values served as the guesses. Upper and lower bounds on the LMO search space were set at 80% and 120% of the originally identified parameter values, respectively, and are listed in the Supplemental Tables. In Table 3 we present the model-fitting statistics for the reduced chi-squared, 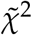, Akaike information criteria (AIC), and Bayesian information criteria (BIC) methods, as well as the parameters determined by LMO. Importantly, we note that fits were performed on data points representing averages and ranges for the two experimental trials at each E:T ratio from only the measured data.

**Table 3.**
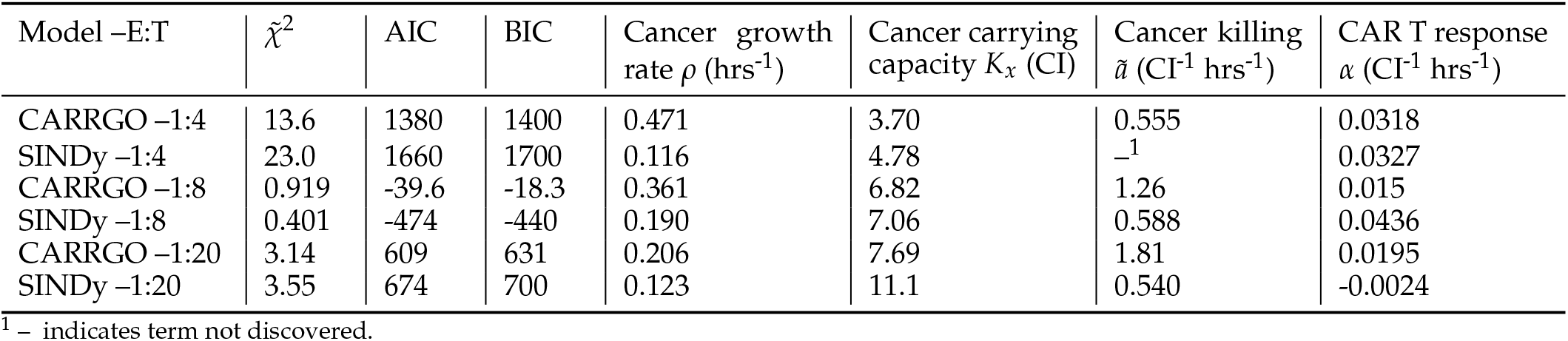
Fitting statistics for CARRGO and SINDy models and comparison of shared parameters. Fitting statistics considered are the reduced chi-squared, 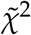, the Akaike information criteria (AIC) and Bayesian information criteria (BIC). Of note are the scores indicating a better fit for the CARRGO model at E:T = 1:4 and 1:20, despite differences in the endpoint CAR T-cell population predictions in Figure 5. Furthermore, we observe generally favorable agreement between parameter estimates, suggesting the *data first* approach of SINDy as a viable alternative to traditional *model first* parameter inference methods.

**Figure 5.**
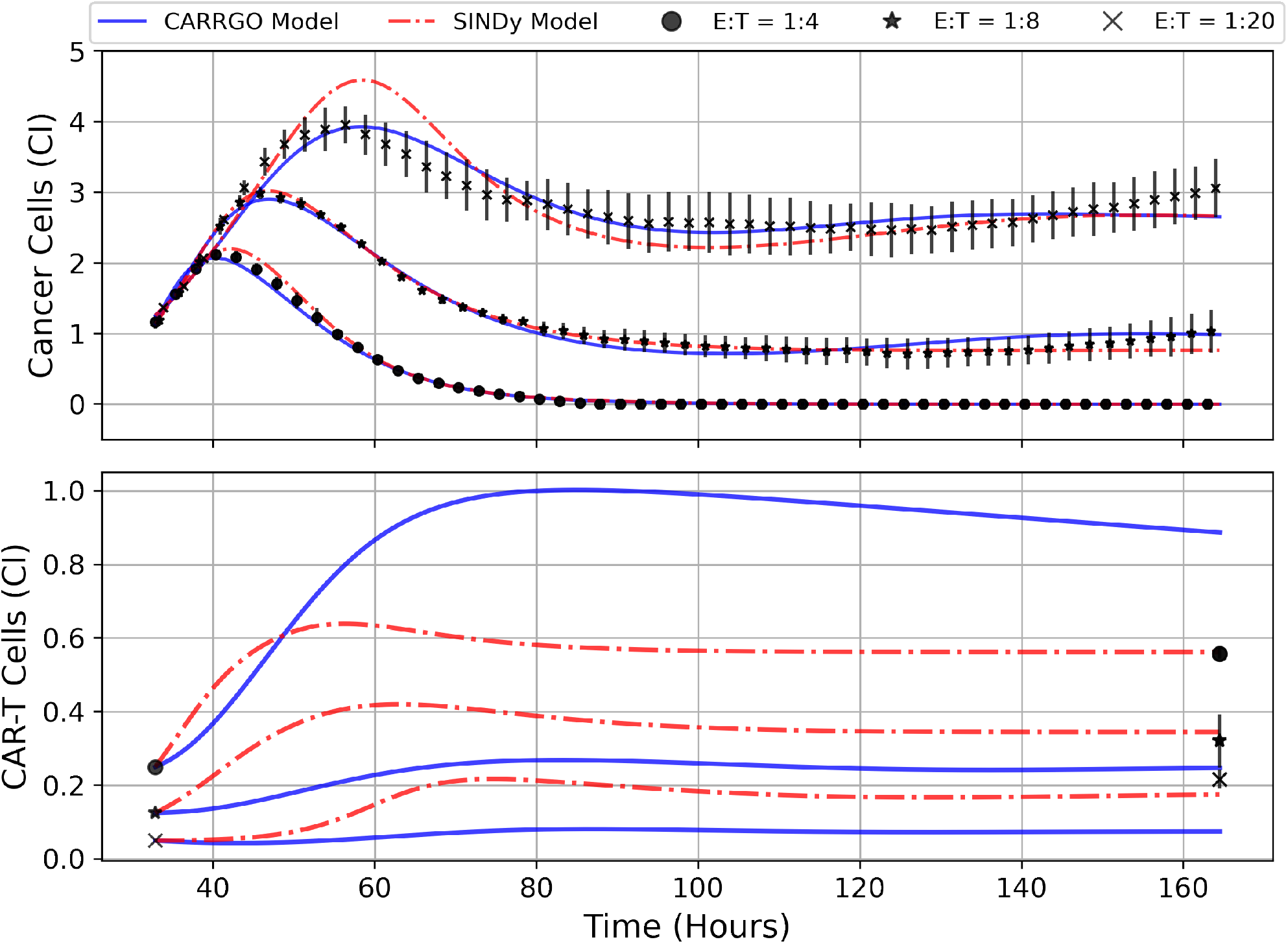
Predictions of cell trajectories for E:T ratios of 1:4, 1:8, and 1:20 from CARRGO model (blue) and SINDy model (red). Model fits for both CARRGO and SINDy were performed using Levenberg-Marquadt Optimization (LMO) on data aggregated across experimental replicates. Initial LMO parameter value guesses were determined by parameter values from SINDy or from published CARRGO model values. Data points represent the mean of all experimental replicates, while error bars represent the ranges across replicates. Of note are the differences in CARRGO and SINDy model predictions for the final CAR T-cell values compared to measurements, and the notable difference in when the maximum CAR T-cell population is reached between CARRGO and SINDy models. Note that experimental measurements have been down-sampled to 25% to allow for visualization.

We find that across the three statistical tests considered, the CARRGO model performs slightly better than the SINDy discovered models at E:T = 1:4 and E:T = 1:20, whereas the SINDy discovered model for E:T = 1:8 performed better than the CARRGO model (Table 3). Interestingly, the CARRGO model predictions for the CAR T-cell trajectories fail to intercept the final CAR T-cell values, where as the SINDy discovered models do. This result highlights a key difference between these two approaches, particularly that the SINDy approach required generating a time-series trajectory for the CAR T-cells that enforced interception with the final CAR T-cell measurement. Alternatively, traditional optimization methods like LMO weight each data point by the range of measurement uncertainty, allowing for the possibility of significant deviation from the final CAR T-cell measurements as long as such deviations can be compensated with better fitting elsewhere amongst the data.

Another essential difference between the CARRGO and SINDy predictions regarding the CAR T-cell trajectories is the CAR T-cell response at the high E:T ratio of E:T = 1:4. Specifically, the CARRGO model predicts that the CAR T-cells reach a maximum population exceeding the maximum population of cancer cells. This result has significant translational implications for CAR T-cell therapy related to patient immune response that we address in the discussion section.

Despite the noted differences, the overall similarities between the CARRGO and SINDy models is demonstrated by the order of magnitude agreement in most shared parameter values, specifically the cancer cell growth rate *ρ*_*x*_, the cancer cell carrying capacity *K*_*x*_, and the CAR T-cell functional response coefficient *α* for the specific scenarios of E:T = 1:4 and E:T = 1:8 (Table 3). Taken together, these results demonstrate significant value in the SINDy methodology when compared to established procedures for parameter estimation.

### 3.3 Model Stability

An important question in performing model discovery for dynamical systems is in relation to the overall stability. Automating the task of examining stability for every discovered model is challenging given the combination of symbolic computation with floating point coefficients. However, by predicting forward in time for each of the models and experimental replicates we can qualitatively characterize the stability (see Figure 4).

For the E:T = 1:4 scenario, both the data and model indicate complete cancer cell death, with the model accurately maintaining a cancer cell population of zero. We note that in several of the alternate discovered models produced by SINDy, the cancer cell population would become negative in the forward predicted regime. This unrealistic result can be used as an aide in ruling out alternative models.

For the E:T = 1:8 and 1:20 scenarios, both the data and models indicate cancer cell-CAR T-cell coexistence, with the forward predictions reaching non-oscillatory steady states. Despite the discovered models being the ones with the best accuracy, they all struggle to match the observed oscillatory frequency, particularly in the E:T = 1:20 scenario. These results demonstrate the capability of SINDy to discover models with variability in solution stability, a core feature of nonlinear dynamical systems.

### 3.4 Parameter Identifiability

To better understand the rarity of the discovered models and their respective coefficients, we examined histograms for the coefficients of each of the model terms along the Pareto fronts for each E:T ratio, presented in Supplementary Figure **S4**. This approach allows us to qualitatively assess parameter identifiability by seeing the extent to which variability in coefficient values exists, and at the expense of prediction accuracy. For most active terms encountered, the coefficients corresponding to the selected models based on the Pareto front analysis were the most commonly occurring values until deactivation (elimination from discovered models). However, in a few situations we see that the coefficient values corresponding to the greatest model accuracy were relatively rare, and varied significantly as increasingly more terms were removed. This occurs in the coefficients for the *x* and *xy*^2^ terms in the cancer cells for the E:T = 1:4 scenario in Supplementary Figure **S4a**, and the *x* and *xy* terms in the cancer cells for both the E:T = 1:8 and 1:20 scenarios in Supplementary Figure **S4b** and **S4c**. These terms were shown to be the final remaining active terms in discovered model, suggesting that they are capable of capturing the greatest extent of variation in our cancer cell-CAR T-cell killing data. Of note once again is that amongst these dominant interaction terms we see a transition from those indicative of double binding at high E:T ratios to single binding at medium and low E:T ratios.

## 4 DISCUSSION

We examined *in vitro* experimental CAR T-cell killing assay data for a human-derived glioblastoma cell line (Figure 1). From our results we infer transitions in the phenomenological killing behavior of the CAR T-cells as a consequence of varying their initial concentration compared to the cancer cells. Our discovered models predict that at high effector to target ratios (E:T = 1:4) the CAR T-cell levels respond according to a Type II functional response in which they survive and/or expand faster at low density, and slower at high density, and they predominantly form double binding conjugates with cancer cells prior to cell killing. At medium E:T ratios of E:T = 1:8 our discovered model again predicts the CAR T-cells undergoing a Type II functional response, but now forming only singly bound conjugates prior to cell killing. At low E:T ratios of E:T = 1:20 our discovered model predicts the CAR T-cells shift to a Type III functional response, in which they survive and/or expand slower at low density, and faster at high density. In this final scenario we find a mixture of single and double conjugate formation occurring. Finally, our discovered models predict the growth strategies of the cancer cells as being a weak Allee effect at high and medium E:T ratios, and logistic at low E:T ratios, while the cancer cells are predicted to follow logistic growth for high and medium E:T ratios. Model coefficients used to deduce these results are found in Tables 1 and 2, and model simulations and forward predictions are shown in Figure 4.

A crucial result of this work is the comparison between the *data first* approach of SINDy to the traditional *model first* approach of CARRGO. Despite the discovered SINDy models having more degrees of freedom (i.e. mathematical terms) than the CARRGO model, both models were found to perform comparably as indicated in Figure 5 and Table 3. Yet, there are key differences regarding the interpretation of these two approaches. Traditional *model first* approaches like the CARRGO model assume a strict individual model that may exhibit variation in its coefficients or model parameters to reflect variation in the underlying biology or experimental conditions. On the other hand, one of the strengths of the *data first* approach of SINDy is that these coefficient variations can be shifted onto discovery of altogether different model terms. As we show, these different terms can have direct interpretations related to the underlying biology and dynamics. For example in (18), variation in the CAR T-cell response due to changes in the experimental E:T ratio could only be indicated through variation in the coefficients of the Type I functional response term, or the value of *α* in Eq. (58). Specifically, increases in *α* were interpreted as a high CAR T-cell response rate, or CAR T-cell expansion, and decreases in *α* were interpreted as a low response rate, or as CAR T-cell exhaustion. Whereas the SINDy model predicts entirely different CAR T-cell functional response terms, providing greater interpretation of these transitions in the CAR T dynamics and biology. Specifically, a Type II functional response at high and medium E:T, or a fast-to-slow CAR T-cell response rate, and a Type III functional response at low E:T, or a slow-to-fast CAR T-cell response that is again suggestive of exhaustion.

### 4.1 Interpreting Discovered Coefficients

We demonstrate the value of the effective model parameters for inferring underlying biology by considering the high E:T model presented in Eqs. (47)-(48). In this scenario, a Type II functional response in the CAR T-cells is deduced from the negative sign on *β*, corresponding to the concave down parabolic nature of the CAR T-cell functional response with fast proliferation at low cancer cell density and slow proliferation at high cancer cell density (Figure 3). The implication that cancer cell killing is induced by double binding of CAR T-cells to cancer cells comes from multiple terms. The most direct indicator is 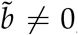, where 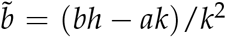 with *bh*/*k* representing the rate of cancer cell death from double conjugates, and *a*/*k* the rate of cancer cell death from single conjugates. Supporting indicators come from the positive sign on *α* = *p*/*g* − *c*/*h*, suggesting that the CAR T-cell death rate from single conjugate formation, *c*/*h* is small compared to the leading order CAR T-cell response rate, *p*/*g*. Further evidence is in the inactivation of the *xy* term in the 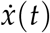 equation with coefficient *ã*. Here, *ã* = *a*/*h* is the rate of cancer cell death from single conjugate formation, whose absence suggests that double binding formation is predominantly responsible for cancer cell death.

A similar analysis of model coefficients for the low and medium E:T ratio scenarios predicts a transition in the interactions between the CAR T-cells and cancer cells. Specifically, our approach predicts that the CAR T-cells form double conjugate pairs with high E:T ratios, then switch to single conjugate pairs at medium and low E:T ratios. Similarly, our results predict a transition in the functional response, indicating Type II functional responses in the CAR T-cells for high E:T ratios and Type III responses in the low E:T ratios. These transitions in detected model terms are phenomenologically consistent with the interactions being dependent on CAR T-cell density, and highlight the hypothesis generating strength of *data first* model discovery techniques. Namely, the prediction of CAR T-cell killing dynamics being dependent on the relative abundance of CAR T-cells compared to cancer cells. We next present several opportunities for experimental testing of these model predictions.

### 4.2 Challenges and Limitations

A challenge to the implementation of SINDy is data sparsity. Despite having high temporal resolution of the cancer cell trajectories (1 measurement per 15 minutes), the CAR T-cell populations consisted of only the initial and final measurements. To resolve sparsity in the CAR T-cell levels, we used latent variable analysis to extract the CAR T-cell trajectory from an approximation to the attractor of the dynamical system as determined by the cancer cell trajectory. We note that in determining the dimensionality of the latent variable subspace, we selected an embedding dimension of *m* = 2 despite the appearance of further benefit in using an embedding dimension of *m* = 3, as indicated in Supplementary Figure **2b**. This choice was made due to our experimental limitations in only having flow cytometry data for the CAR T-cells at the initial and final time points, and no further data with which to constrain any additional latent variables. The existence of a second latent variable, as suggested by the third embedding dimension, could be due to single or double binding conjugates if the reaction rates are sufficiently slow, or, alternatively, a biochemical secretion that is modulating the cancer cell and CAR T-cell interactions. Future experimental and modeling efforts may further illuminate the nature of this third state variable, which we discuss in the Future Directions section.

One potential limitation with latent variable analysis is that the trajectories retrieved through Taken’s Theorem are not guaranteed to be unique, but rather will be diffeomorphic to the true latent variable. That is, subject to topological stretching or skewing, which translates to variation in discovered model coefficients. This effect can be seen in Bakarji et. al (62), where the coefficients of the latent variables discovered for the two-state, predator-prey model are not in precise agreement to those used in the original simulation. However, it is important to note that the model terms discovered by SINDy with this methodology are biologically insightful, even though the coefficients multiplying the discovered model terms on latent variables may be subject to variation. Importantly, we provide further experimental information for the latent CAR-T cell variable through bounding of the initial and final CAR-T cell trajectory with direct measurements. Likewise, we only discover terms which are structurally identifiable through model inversion, minimizing the potential for diffeomorphic skewing of CAR-T cell trajectories to be discovered from Taken’s Theorem.

A second challenge is that our data in total consists of two trials for each effector-to-target ratio. While there exist SINDy implementations designed to discover models with ratios of polynomials, the approaches require prohibitively many experimental trials to ensure accuracy (39, 40). To resolve sparsity in the number of experimental trials, we derived effective interaction models of cancer cell and CAR T-cell dynamics from model ODE terms with ratios of the polynomials using binomial approximations. These effective interaction models allowed for the identification of multiple constraints on the library function space used in SINDy, and guided our inferential analysis of the discovered models.

### 4.3 Future Directions and Clinical Applications

To validate the hypothesized binding and functional response dynamics, we propose two potential experiments. Both experiments rely on similar initial conditions as those conducted for this study, but in one we propose the use of bright field microscopy and live cell imaging to visually inspect CAR T-cell dynamics at different points in time and for the different E:T ratios. By tracking in real-time the growth, motility, and interactions of the different cells present, this approach ought to aide in distinguishing different cell phenotypes by identifying occurrences of single and double binding types as well as the different functional responses (63). The second experiment would be to conduct endpoint analyses using flow cytometry to determine the population of CAR T-cells throughout the trajectory. This experiment would test the different CAR T-cell predictions from the CARRGO model and the SINDy models, most notably the predicted time to reach maximum CAR T-cell populations (Figure 5). Furthermore, targeted staining can provide information on the number of CAR T-cell generations and the ratio of helper T-cells (CD4+) to cytotoxic, killer T-cells (CD8+). These metrics may better inform the number of true effector cells responsible for killing cancer cells, allowing for more accurate characterization of the CAR T-cell response. These experiments additionally serve to test the validity of our latent variable analysis, which uses the cancer cell trajectory to predict the CAR T-cell trajectory as presented in Figure 4. Future experiments will also extend this analysis to include other CAR designs, including evaluating the impact of costimulatory signaling, CAR affinity and target density on modeling of CAR T-cell killing dynamics.

These and other experiments are essential for introducing additional elements and agents present in the tumor microenvironment and for extending this work to *in vivo* applications. Currently, our implementation of SINDy is on a highly controlled experimental system in order to isolate the interaction dynamics between the CAR T-cells and the glioma cells and to validate the SINDy methodology. An important challenge to overcome is extending the SINDy framework to incorporate additional aspects of *in vivo* systems. To achieve this, intermediate experiments to conduct are killing assays in two- and three-dimensional *in vitro* tissue model systems that mimic the tumor microenvironment (64). The proposed experiments are crucial for adapting use of the SINDy framework for clinical applications.

The clinical relevance of the *data first* framework is in the domain of precision medicine. The approach naturally caters to *in situ* monitoring of patient response to therapy and forecasting future trajectories. An open question in this field is determining the sufficient number of early measurements necessary for accurate forecasting, and quantifying the extent of reliable forward prediction. This type of application falls under the field of control theory, in which real-time measurements for systems such as navigation, fluid dynamics and disease monitoring can inform model-based interventions (15). Control theory has been identified as a key tool in achieving optimized individual treatment outcomes, yet challenges are ever-present in parsimonious model selection. The SINDy methodology may help streamline and simplify the model selection process, while simultaneously incorporating control theory methods for treatment optimization. As an example related to the experiments considered here, one could envision a therapeutic intervention to administer more CAR T-cells in the low E:T ratio of 1:20 as soon as the Type III functional response and single binding dynamics are predicted in a patient. This intervention would serve to push the dynamics of the patients immune response into the double biding and Type II response regime, thereby improving therapeutic efficacy.

## 5 CONCLUSIONS

In this work we present the first, to our knowledge, application of the sparse identification of non-linear dynamics (SINDy) methodology to a real biological system. We used SINDy with highly time-resolved experimental data to discover biological mechanisms underlying CAR T-cell-cancer cell killing dynamics. Our implementation highlights the hypothesis generating potential of data-driven model discovery and illuminates challenges for future extensions and applications. To overcome challenges related to data limitation, we utilized latent variable analysis to construct the trajectory of the CAR T-cells, and we implemented binomial expansions to simplify specific model terms. Our results predict key mechanisms and transitions in the interaction dynamics between the CAR T-cells and cancer cells under different experimental conditions that may be encountered in the application of these therapies in human patients. Specifically, we identified transitions from double CAR T-cell binding to single CAR T-cell binding, and from fast-to-slow CAR T-cell responses (Type II) to slow-to-fast responses (Type III). Both transitions occur as a result of decreasing the relative abundances of CAR T-cells to cancer cells (initial E:T ratios). Importantly, these results demonstrate the potential for *data first* model discovery methods to provide deeper insight into the underlying dynamics and biology than *model first* approaches, and offer a new avenue for integrating predictive modeling into precision medicine and cancer therapy by an improved mechanistic understanding of cancer progression and efficacy of CAR T-cell therapy.

## AUTHOR CONTRIBUTIONS

Conceptualization, AB and RR; methodology, AB, RW, VA, HC and RR; validation, AB, RR; formal analysis, AB; investigation, AB; resources, MG, CB, and RR; data curation, AB; writing— original draft preparation, AB; writing—review and editing, AB, AX, AW, VA, HC, MG, CB, RR; visualization, AB, AX, RW, VA, RR; supervision, CB and RR; project administration, AB and RR; funding acquisition, CB and RR All authors have read and agreed to the published version of the manuscript.

## SUPPLEMENTAL MATERIALS

Supplemental Table 1 - Parameter seed values for Levenberg-Marquardt Optimization of CARRGO and SINDy growth-death model terms.

Supplemental Table 2 - Parameter seed values for Levenberg-Marquardt Optimization of CARRGO and SINDy interaction model terms.

Supplemental Figure 1 - Determination of latent variable analysis time delay and embedding dimension values.

Supplemental Figure 2 - Latent variable analysis results and SINDy predicted models for second replicate of experimental conditions.

Supplemental Figure 3 - Pareto front analyses investigating trade off in model complexity and model accuracy.

Supplemental Figure 4 - Examination of frequency of discovered terms and associated coefficients as threshold *λ* is varied in model discovery.

## DATA AVAILABILITY STATEMENT

The data presented in this study and Python code used to analyze data and generate figures are openly available from the following online location https://github.com/alexbbrummer/CART_SINDy.

