## Supplemental Material for "Data driven model discovery and interpretation for CAR T-cell killing using sparse identification and latent variables"

### 1 SUPPLEMENTARY TABLES

**Table S1.** Parameter seed values for Levenberg-Marquardt Optimization of CARRGO and SINDy growth-death model terms. Upper and lower bounds were set a 20% above and below seed values for all present terms.

| Model –E:T | $\rho_x$ (hrs$^{-1}$) | $A_x/B_x$ (CI) | $K_x$ (CI) | $\rho_y/\theta_y$ (hrs$^{-1}$) | $A_x/B_x$ (CI) | $K_y$ (CI) |
|---|---|---|---|---|---|---|
| CARRGO –1:4 | 0.471 | –[1] | 3.70 | 0.00182 | – | – |
| CARRGO –1:8 | 0.359 | – | 5.69 | 0.0134 | – | – |
| CARRGO –1:20 | 0.229 | – | 6.15 | 0.0645 | – | – |
| SINDy –1:4 | 0.121 | 1.42 | 4.79 | 0.191 | – | 0.544 |
| SINDy –1:8 | 0.237 | 3.08 | 6.41 | 0.112 | – | 0.313 |
| SINDy –1:20 | 0.15 | – | 12.5 | – | – | – |

[1] – indicates term not a part of model.

**Table S2.** Parameter seed values for Levenberg-Marquardt Optimization of CARRGO and SINDy interaction model terms. Upper and lower bounds were set a 20% above and below seed values for all present terms.

| Model –E:T | $\tilde{a}$ (CI$^{-1}$ hrs$^{-1}$) | $\tilde{b}$ (CI$^{-2}$ hrs$^{-1}$) | $\alpha$ (CI$^{-1}$)(hrs$^{-1}$) | $\beta$ (CI$^{-1}$ hrs$^{-1}$) | $\tilde{c}$ (CI$^{-2}$ hrs$^{-1}$) |
|---|---|---|---|---|---|
| CARRGO –1:4 | 0.555 | –[1] | 0.0319 | – | – |
| CARRGO –1:8 | 1.045 | – | 0.0159 | – | – |
| CARRGO –1:20 | 1.51 | – | 0.0244 | – | – |
| SINDy –1:4 | – | 0.593 | 0.035 | -0.009 | – |
| SINDy –1:8 | 0.626 | – | 0.051 | -0.0096 | – |
| SINDy –1:20 | 0.545 | – | -0.002 | 0.005 | 0.063 |

[1] – indicates term not discovered.

### 2 SUPPLEMENTARY FIGURES

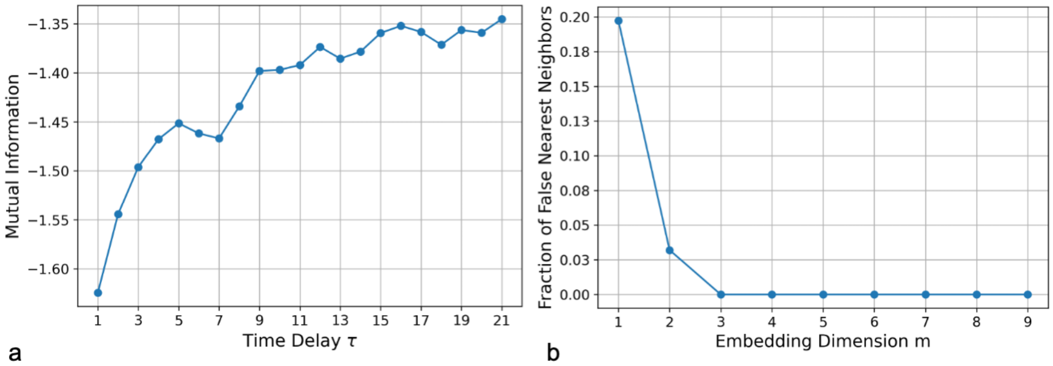


a b

**Figure S1.** Graphs for determining **a** the ideal time delay, $\tau$, by examining mutual information and **b** the ideal embedding dimension, $m$, by examining the fraction of false nearest neighbors. For our data, the optimal time delay found was $\tau = 1$ and the optimal embedding dimension $m = 2$.

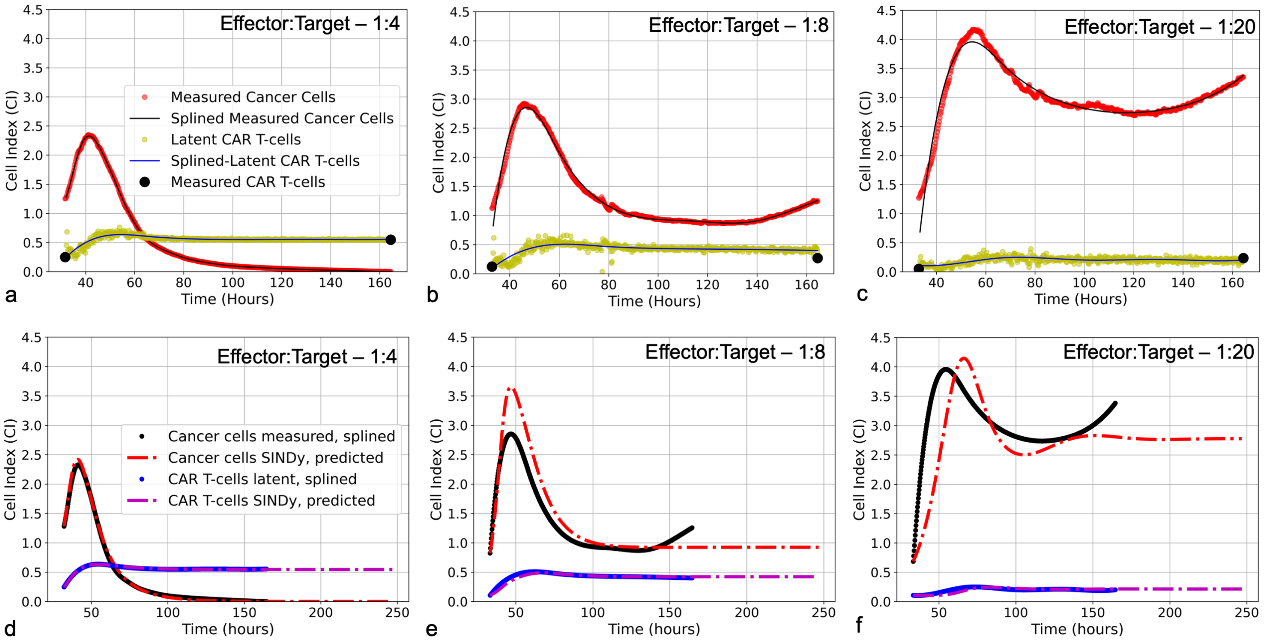


**Figure S2.** (**a-c**) Latent variable analysis results for second of two experimental replicates for each E:T ratio examined. Presented for are the cancer cell index measurements from the xCELLigence machine in red, overlaid with the splined measurements for the cancer cells in black; the two endpoint measurements for the CAR T-cell levels enumerated by flow cytometry in black, with the CAR T-cell population trajectory as determined by latent variable analysis in yellow, overlaid with the splined CAR T-cell trajectory in blue. Note that despite the CAR T-cell populations being measured with flow cytometry, we have converted levels to units of Cell Index for ease of comparison with the cancer cells, using a conversion factor of $1\text{CI} \approx 10,000$ cells. (**d-f**) Predicted trajectories of discovered models compared to splined measurements of cancer cells and CAR T-cells for same data presented in (**a-c**). Splined cancer cell and CAR T-cell measurements are in black and blue, respectively. Predicted trajectories for cancer cells are the red dot-dashed lines, while the CAR T-cells are the purple dot-dashed lines. To examine stability of SINDy-discovered models, both simulations and forward predictions are presented to show steady-state behavior. Note that the best fits between predictions and measurements occur in the high E:T scenario, where assumptions made regarding treatment success and low cancer cell populations in determining model candidate terms are best adhered. As the E:T ratios get smaller, increasing deviation between discovered model predictions and splined measurements can be qualitatively observed. This is likely due to weakening of assumptions of treatment success and low cancer cell populations associated with the low E:T conditions.

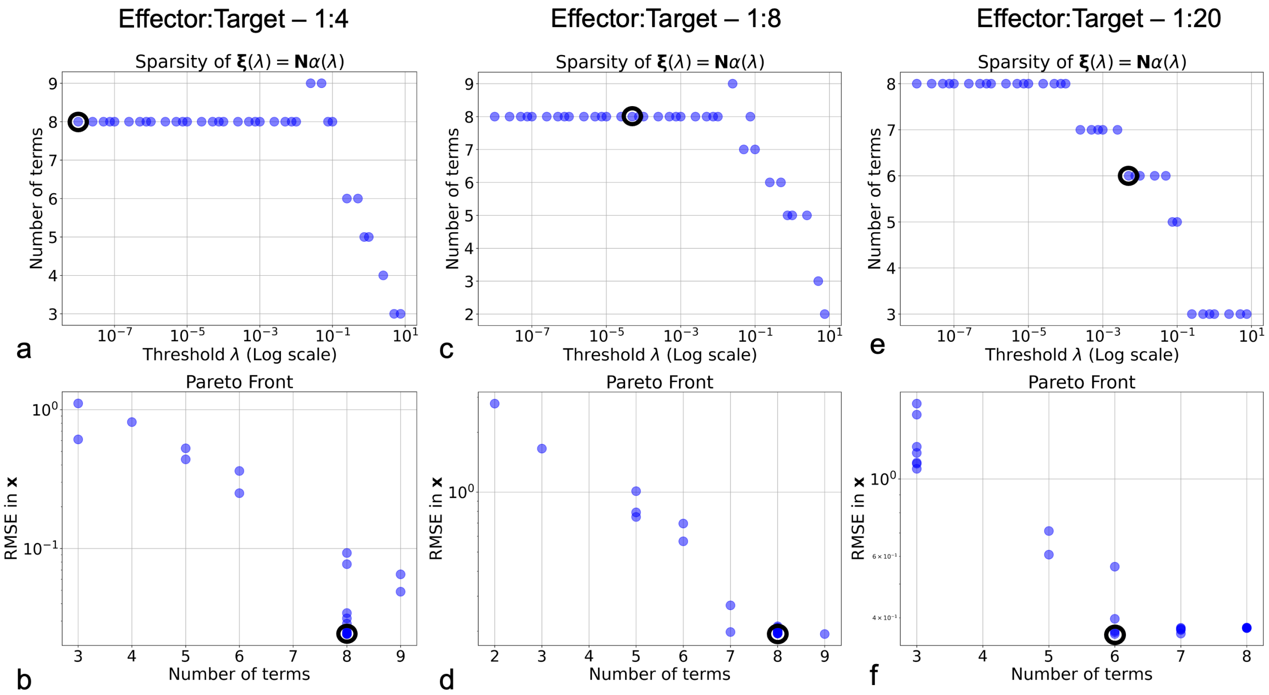


**Figure S3.** Pareto front analysis of system sparsity for the low, medium and high effector:target ratios (E:T = 1:20, E:T = 1:8 and E:T = 1:4). **a, b, c** present model sparsity (number of discovered terms) versus threshold $\lambda$ and **d, e, f** present root-mean-squared-error (RMSE) between SINDy-model predictions and measurements versus number of terms. Note that the RMSE values were calculated using the discovered models and the splined measurements. Selected models are represented by purple circles.

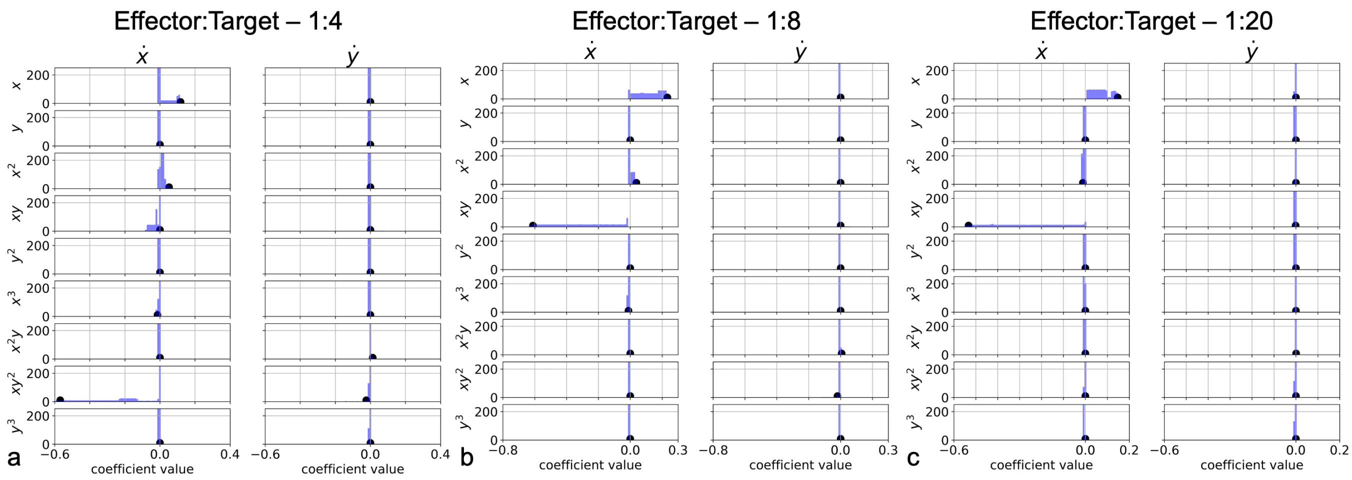


**Figure S4.** Frequency of discovery for model terms as threshold $\lambda$ is varied across 1000 values from the interval $[5^{-3}, 10^{1}]$ for **a** high E:T, **b** medium E:T, and **c** low E:T. Black circles indicate values for coefficients corresponding to the selected model based on the Pareto front analysis in Figure S3.
